# Enteropathogenic *E. coli-*mediated Fast and Coordinated Ca^2+^ responses regulate NF-κB activation

**DOI:** 10.1101/2025.08.06.668902

**Authors:** Fangrui Guo, Roberto Ornelas Guevara, Linda Oussaedine, Geneviève Dupont, Laurent Combettes, Guy Tran Van Nhieu

## Abstract

Enteropathogenic *Escherichia coli* (EPEC) is a major bacterial enteropathogen causing infectious diarrhea among children in developing countries. Here, we found that EPEC induced isolated Ca^2+^ responses in epithelial cells, triggered by extracellular ATP (eATP). These responses were dependent on type III secretion (T3S) and down-regulated by the bacterial secreted protease EspC, consistent with eATP released by the T3S translocon pore-forming activity in host membranes. By performing high speed Ca^2+^ imaging, we uncovered that at the onset of infection, low eATP levels triggered Ca^2+^-responses involving the whole cell but showing the small amplitude and fast kinetics usually associated with local Ca^2+^ responses. The findings, supported by theoretical modeling, evocate a conceptual shift whereby low amounts of inositol 1, 4, 5-trisphosphate (IP_3_) induced by low eATP levels and subsequent moderate Ca^2+^ release enable the fast coordination of IP_3_ receptor cluster activation throughout the cell. Importantly, these yet undescribed coordinated fast responses occurred over prolonged time periods and defined a cell state with dampened activation of the pro-inflammatory transcriptional activator NF-kB associated with a decrease in its Ca^2+^-dependent O-linked β-N-acetylglucosamine modification.

## Introduction

EPEC are diarrheagenic *E. coli* strains that cause significant morbidity and mortality in children under two years of age. While infection rates have significantly declined in industrialized nations, EPEC remains a major public health concern in low-income countries (Lozer et al., 2013). EPEC form attaching and effacing (A/E) lesions on intestinal epithelial cells and lack the ability to produce Shiga toxins or heat-labile (LT) and heat-stable (ST) enterotoxins (Croxen et al., 2013; Gomes et al., 2016; Hazen et al., 2016). The ability of EPEC to form A/E lesions is determined by the locus of Enterocyte Effacement (LEE), a large genomic pathogenicity island that encodes the essential genetic elements required for this process (Pearson et al., 2016). The LEE region of EPEC (E2348/69) encodes components of the type III secretion system (T3SS), a molecular apparatus that translocate at least 25 bacterial effector proteins into the host cell. The EPEC type III secretion system (T3SS) consists of a basal body and a needle-like structure, resembling those found in *Salmonella* and *Shigella*, but with a distinct sheath-like extension at the needle tip, primarily composed of EspA, and about ten times longer than the T3SS needles from other bacterial species. This EspA filament acts as a molecular bridge, extending from the bacterium to the host cell membrane, allowing insertion of the EspB and EspD translocon components into the host cell membrane, enabling type III effectors injection in the cell cytosol (Creasey et al., 2003; Monjarás Feria et al., 2012; Sal-Man et al., 2012). Osmoprotection assays suggest that the translocon forms a pore with an internal diameter ranging between 3 to 5 nm, allowing the passage of unfolded effector proteins into the host cell (Chatterjee et al., 2015). However, the T3SS translocon shows weak pore-forming activity during EPEC infection of epithelial cells, presumably because it forms a sealed conduct between the T3SS and host cell membranes (Guignot et al., 2016). EspC, a secreted serine protease from the autotransporter family, targets EspA and EspD and down-regulates pore formation activity associated with cytotoxicity (Guignot et al., 2015). EPEC T3SS effector proteins translocated into host cells lead are responsible for attaching and effacing (A/E) lesions and intimate bacterial adhesion to the host cells associated with the formation of an actin-rich pedestal formation (Chen & Frankel, 2005).

The detection of pathogenic bacteria by intestinal epithelial cells plays an important role in initiating pro-inflammatory responses. Recognition of bacterial surface components by pattern recognition receptors triggers pro-inflammatory signaling pathways involving the transcriptional activator NF-κB and the production of cytokines such as interleukin-8 and tumor necrosis factor-α (TNF-α) (Edwards et al., 2011). However, EPEC suppresses these signaling pathways early in infection through the coordinated action of several T3SS effector proteins. Among the first translocated T3SS effectors, Tir interacts with TNF-α receptor-associated factors (TRAF2 and TRAF6), recruiting the tyrosine phosphatases SHP-1 and SHP-2 (Mills et al., 2008; Ruchaud-Sparagano et al., 2011; Yan et al., 2013). Several non-LEE T3SS effectors, including NleE, NleB, NleH1, NleH2, NleC, and NleD, further contribute to inhibiting NF-κB and MAPK signaling. NleE and NleB stabilize the interaction between NFκB and its inhibitory subunit IκB, preventing its degradation, thereby keeping NFκB in an inactive state. The ability of NleE to inhibit NFκB signaling depends on its S-adenosyl-L-methionine (SAM)-dependent methyltransferase activity (Zhang et al., 2011). NleE-mediated methylation of TAB2/3 prevents IKK activation (Zhang et al., 2011). NleB selectively blocks NFκB activation through GlcNAcylation of the TNF-α receptor (TNFR) adaptor protein and TNFR1-associated death domain (TRADD) (S. Li et al., 2013; Pearson et al., 2013). The T3SS effectors NleH1, NleH2 and NleC also interfere with NFκB nuclear translocation (Gao et al., 2009). NleC specifically cleaves P65 RelA (Giogha et al., 2015; Ruchaud-Sparagano et al., 2011; Yen et al., 2010). Interestingly, NleF has been implicated in the activation of NF-κB, underscoring the complexity of the regulation of inflammation during bacterial infection and suggesting the timing of various T3SS effectors’ activity (Pallett et al., 2014).

A similar complexity applies to T3SS effectors regulating cell death and survival pathways during EPEC infection of epithelial cells. Tir was found to elicit a rapid Ca²⁺ influx across the host cell membrane, through the activation of a host plasma membrane Ca²⁺ channel, the mechanosensitive transient receptor potential vanilloid 2 (TRPV2) leading to pyroptosis (Zhong et al., 2022). However, the NleA effector, blocks the delivery of TRPV2 channels to the cell surface, thereby dampening Tir-induced Ca²⁺ influx. The effector NleF also directly binds caspase-4 to inhibit its activity (Zhong et al., 2020). The extrinsic apoptotic pathway is triggered by EPEC pili (Abul-Milh et al., 2001). However, the NleD and NleB effectors inhibit this pathway by cleaving JNK and GlcNAcylating the death domain adaptor proteins TRADD and FADD, respectively (Baruch, Gur-Arie, et al., 2011; Pearson et al., 2013). While EspC prevents cytotoxicity linked to pore-formation by the T3SS translocon during the early EPEC infection phases, it was shown to promote intrinsic apoptosis through increase in intracellular Ca^2+^ and calpain activation (Serapio-Palacios & Navarro-Garcia, 2016).

Central to inflammation and cell death / survival pathways induced by EPEC, bacterial-induced Ca^2+^ signals have been a matter of debate. EPEC infection is known to perturb host Ca²⁺ signaling, but the source and sequence of Ca²⁺ signals during infection remain controversial. EPEC was shown to induce Ca²⁺ influx associated with a loss of mitochondrial membranes permeability leading to cell death (Zhong et al., 2020; Ramachandran et al., 2020; Zhong et al., 2022), but was also reported to trigger IP_3_-mediated Ca²⁺ release possibly involved in bacterial-induced cytoskeletal rearrangements (Baldwin et al., 1991; Baldwin et al., 1993; Foubister et al., 1994; Bain et al., 1998).

Here, we investigated the characteristics and implications of EPEC-induced Ca²⁺ responses in epithelial cells. We characterized yet undescribed Ca²⁺ signals induced by EPEC and low ATP levels, presenting the fast dynamics and small amplitude of local Ca²⁺ responses but involving large cell area. We found that these responses likely result from the coordination of elementary responses via rapid Ca^2+^-induced Ca²⁺ release over large cell area, challenging generally admitted concepts on Ca²⁺ diffusion. Importantly, we show that these newly described responses have functional implications by dampening the cell ability to respond to inflammatory signals.

## Results

### EPEC induces Ca^2+^ responses that depend on Type III secretion-mediated eATP release

Despite their critical role in cellular processes key to bacterial infection, EPEC-induced Ca²⁺ responses remain to be characterized. We therefore set up to perform a detailed single cell imaging of Ca²⁺ responses elicited by cells infected by EPEC.

As shown in Fig. 1, EPEC induced Ca²⁺ transients often corresponding to a single peak of varying amplitude detected over several minutes, corresponding to 6.2 ± 0.8% (mean ± SEM) of the maximal histamine response (Figs. 1A, B). These Ca²⁺ responses were dependent on a functional T3SS, since they were not observed for the T3SS-deficient *escN* mutant (Figs. 1A, C). Only 40 ± 4.7 % (mean ± SEM) of cells, however, elicited responses when challenged wild-type EPEC at a low multiplicity of infection (MOI) of 20 bacteria per cell, a value that raised to 76 ± 4.8 % (mean ± SEM) when using a high MOI of 80 bacteria per cell (Fig. 1C). In contrast, even at the low MOI, more than 83 % of cells showed actin pedestals, indicating that Ca²⁺ responses were elicited only in a fraction of cells targeted by EPEC-type T3SS (Figs. 1D, E). The frequency of Ca²⁺ responses per cell increased over the incubation time with an average frequency of responses per cell raising by 6.9 to 8.3-fold from the first to the last 30 min of EPEC challenge (Fig. 1F). This increased frequency suggested the accumulation of an agonist in the extracellular medium during the course of the infection triggering IP_3_-mediated Ca²⁺ release. When pooling all single cell responses, we could observe a steady increase in the average cytosolic Ca^2+^ concentration of the cell population, as previously reported (Ramachandran et al., 2020; Figs. S1B). However, when performing single cell imaging, we did not observe an increase in cytosolic Ca²⁺ basal levels even at high MOI after 2 hours incubation with wild-type bacteria (Fig. S1A).

**Figure 1.**
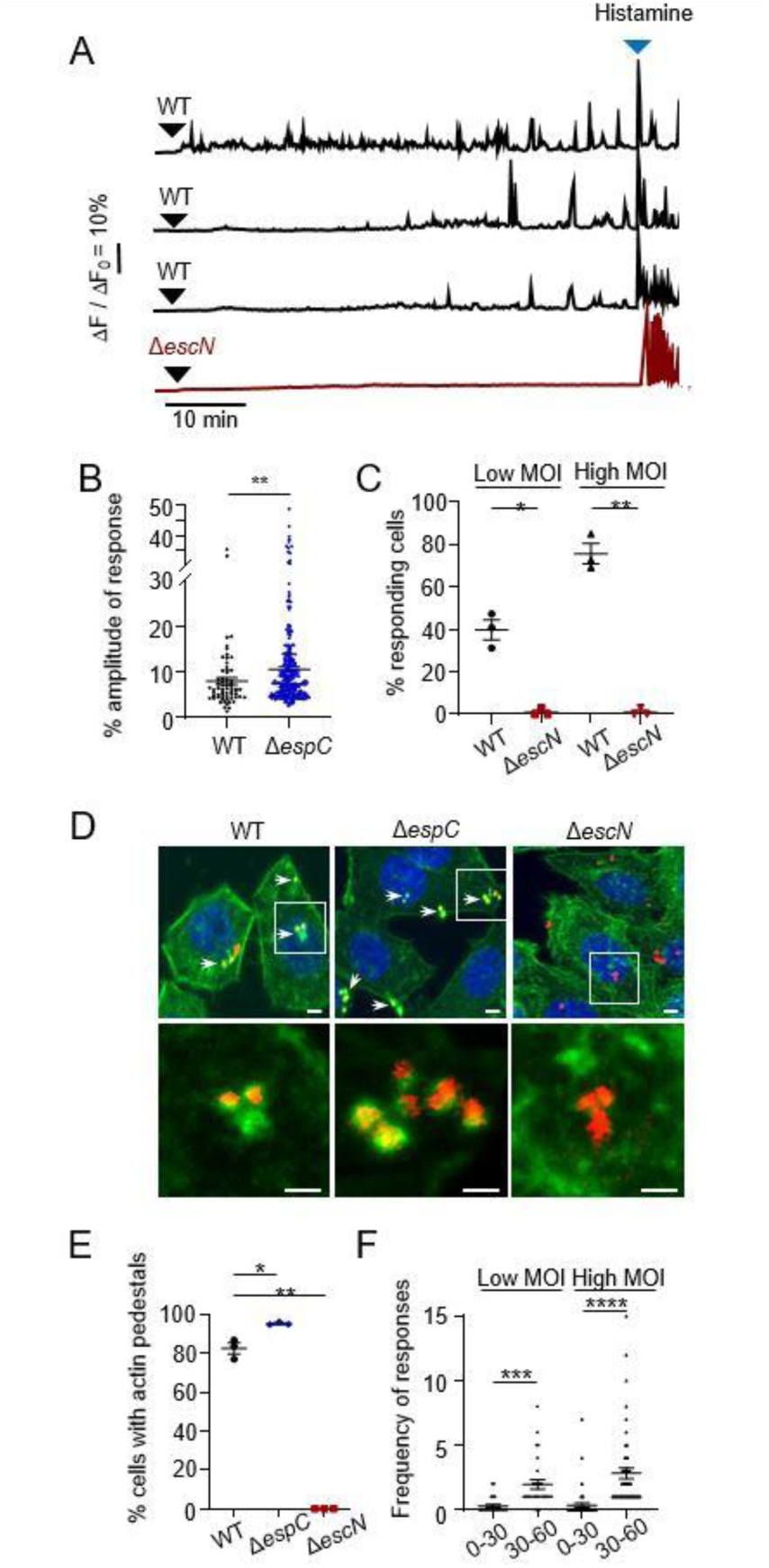
EPEC induces isolated Ca^2+^ responses of limited amplitude in epithelial cells. HeLa cells were loaded with the fluorescent indicator Cal-520, challenged with the indicated bacteria and subjected to live-cell Ca^2+^ imaging at a frequency of one acquisition every 10 seconds (**A-C, F**) or fixed and processed for fluorescence microscopy analysis (**D-E**) (Materials and Methods). **A**, Representative traces of Ca^2+^ variations in single cells. The black arrowheads indicate the time of bacterial challenge. The blue arrowheads indicate stimulation with 3 µM histamine. **B**, Response amplitude expressed as a percent of the maximal histamine response amplitude (N = 3, n > 63). **C**, Percent of cells exhibiting Ca^2+^ responses (N =3, cells > 66). (**D, E**) Cells challenged with RFP-expressing bacteria for 1 hour. **D**, Representative confocal micrographs. Staining with DAPI (blue), phalloidin-Alexa 488 (green). The lower panels show a higher magnification of the insets in the top panels. Scale bar = 10 µm. **E**, Percentage of bacteria-associated actin-rich pedestals (N = 3, cells > 273). **F**, average number of responses per cell during the first 30 min (0-30) and last 30 min (30-60) of bacterial challenge. Low MOI: 10 bacteria / cell. High MOI: 50 bacteria / cell. Bar: mean. (N = 3, cells > 63). Mann-Whitney test. *: p < 0.05; **: p < 0.01; ***: p < 0.001; ****: p < 0.0001.

Together, these results suggest that EPEC induces isolated Ca²⁺ responses that depend on the T3SS for the release of limiting amounts of a Ca²⁺ agonist.

### EPEC-mediated Ca²⁺ responses depend on ATP released in the extracellular medium via the T3SS translocon

In previous works, we showed that the EPEC T3SS translocon forms pores in host cell plasma membranes that were down-regulated by the bacterial secreted serine protease EspC (Guignot et al., 2016). We posit that low amounts of ATP released in the extracellular medium by the T3SS translocon were responsible for the isolated Ca²⁺ responses of reduced amplitude elicited by EPEC. According to this view, by removing T3SS translocons from host cell membranes, EspC would down-regulate EPEC-mediated Ca^2+^ signaling explaining the low ratio of Ca^2+^ responding cells relative to cells forming actin pedestals.

Consistent with this and as shown in Figs. 2A-C, an *espC* mutant induced more Ca²⁺ responses than wild-type EPEC, with 94 ± 3% responding cells and a frequency of 10.5 ± 1.2 responses per cell over the 60 min analysis, compared 40 ± 4.7% responding cells and less than 2 responses per cell for the *espC* mutant and wild-type EPEC, respectively. Also, the average amplitude of Ca²⁺ responses induced by the *espC* mutant was higher than that of wild-type EPEC, suggesting more eATP release (Fig. 1B). Accordingly, cell treatment with the purinergic receptors’s antagonists Suramin and PPADS as well as with hexokinase to deplete eATP, inhibited Ca²⁺ responses induced by the wild-type and *espC* mutant strains (Figs. 2B, C, S2A-C). Treatment with the PLC inhibitor U73122, but not its inactive analog U73343, also resulted in inhibition of EPEC-mediated Ca²⁺ responses (Figs. S2A, B). As expected for ATP-mediated Ca^2+^ release, sample treatment with EGTA, a cell impermeant chelator of extracellular Ca^2+^ did not decrease the percent of Ca^2+^ responding cells triggered by wild-type EPEC or the *espC* mutant (Fig. S2C). The frequency of responses per cell was also not inhibited and even appear to increase upon EGTA-treatment in cells challenged with wild-type EPEC and the *espC* mutant (Figs. S2D, E). In control experiments, Suramin treatment did not affect actin pedestal structures induced by these strains (Fig. S3).

**Figure 2.**
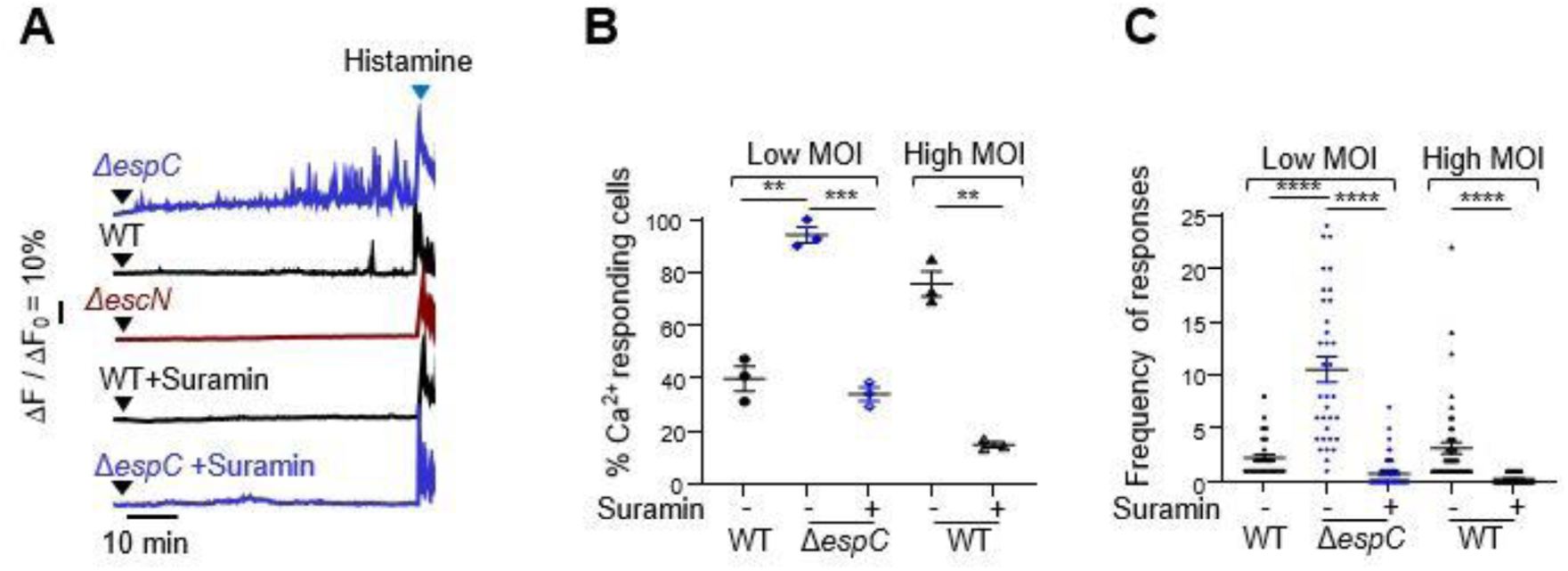
EPEC-induced Ca^2+^ responses are elicited by ATP released by the T3SS translocon. HeLa cells were loaded with the fluorescent indicator Cal-520 or with 200 μM suramin for 30 minutes, challenged with the indicated bacteria and subjected to live-cell Ca^2+^ imaging for a 60 min-duration at a frequency of one acquisition every 10 seconds. **A**, Representative traces of Ca^2+^ variations in single cells. The black arrowheads indicate the time of bacterial challenge. The blue arrowheads indicate stimulation with 3 µM histamine. **B**, Percent of cells exhibiting Ca^2+^ responses (N = 3, cells > 70). **C**, Average number of responses per cell. Low MOI: 10 bacteria / cell. High MOI: 50 bacteria / cell. Bar: mean. (N = 3, cells > 30). Mann-Whitney test. **: p < 0.01; ***: p < 0.001; ****: p < 0.0001.

Together, these results suggest that Ca²⁺ responses induced by EPEC are mediated by ATP released in the extracellular medium via pores formed by the T3SS translocon and are down-regulated by EspC.

### EPEC induces coordinated Ca²⁺ responses from single IP_3_R clusters

We previously showed that *Shigella* induced local Ca^2+^ responses dependent on the T3SS and Ca^2+^ release (Tran Van Nhieu et al., 2013), suggesting that insertion of the Type III translocon was responsible for bacterial-induced local Ca^2+^ signals. We therefore set up to investigate whether EPEC could also trigger T3SS-dependent local Ca^2+^ responses.

To explore this, we performed rapid Ca^2+^ imaging at a frequency of 57 ms acquisition per frame to sample elementary Ca^2+^ release events. As shown in Fig. 3, by performing high speed Ca^2+^ imaging, we detected fast Ca^2+^ increases associated with cell challenge with EPEC. These fast Ca^2+^ responses did not correspond to other responses previously reported since they involved the whole cell or a large cell area but showed a small amplitude and fast dynamics usually associated with local Ca^2+^ responses (Fig. 3B). As observed for the ATP-dependent responses shown in Fig. 2, the *espC* mutant triggered a higher percent of Ca^2+^ responding cells than wild-type EPEC (Fig. 3C). Also, the response amplitude was higher for the *espC* mutant with an average amplitude corresponding to 7.7 ± 0.4 % (mean ± SEM) of the maximal agonist response, compared to 5.4 ± 0.4 % (mean ± SEM) for wild-type EPEC (Fig. 3E). These responses occurred repeatedly at a high frequency of up to 4.5 responses per minute during several minutes following bacterial challenge (Figs. 3D and 3F) and were dependent on Type III Secretion, as evidenced by the lack of response in cells infected with the *ΔescN* strain (Figs. 3B and 3C). EPEC-induced fast Ca^2+^ responses were dependent on Ca^2+^ release since they were inhibited by U73122, a PLC inhibitor (Figs. 3C and 3D). These similarities with the EPEC-induced eATP-dependent Ca^2+^ responses suggested that the atypical fast responses were triggered by low amounts of ATP released in the extracellular medium following insertion in host cell plasma membranes of discrete numbers of EPEC T3SS translocons. Consistently, these atypical EPEC-induced fast responses were abolished in the presence of the ATP receptor inhibitor Suramin (Figs. 3C, D).

**Figure 3.**
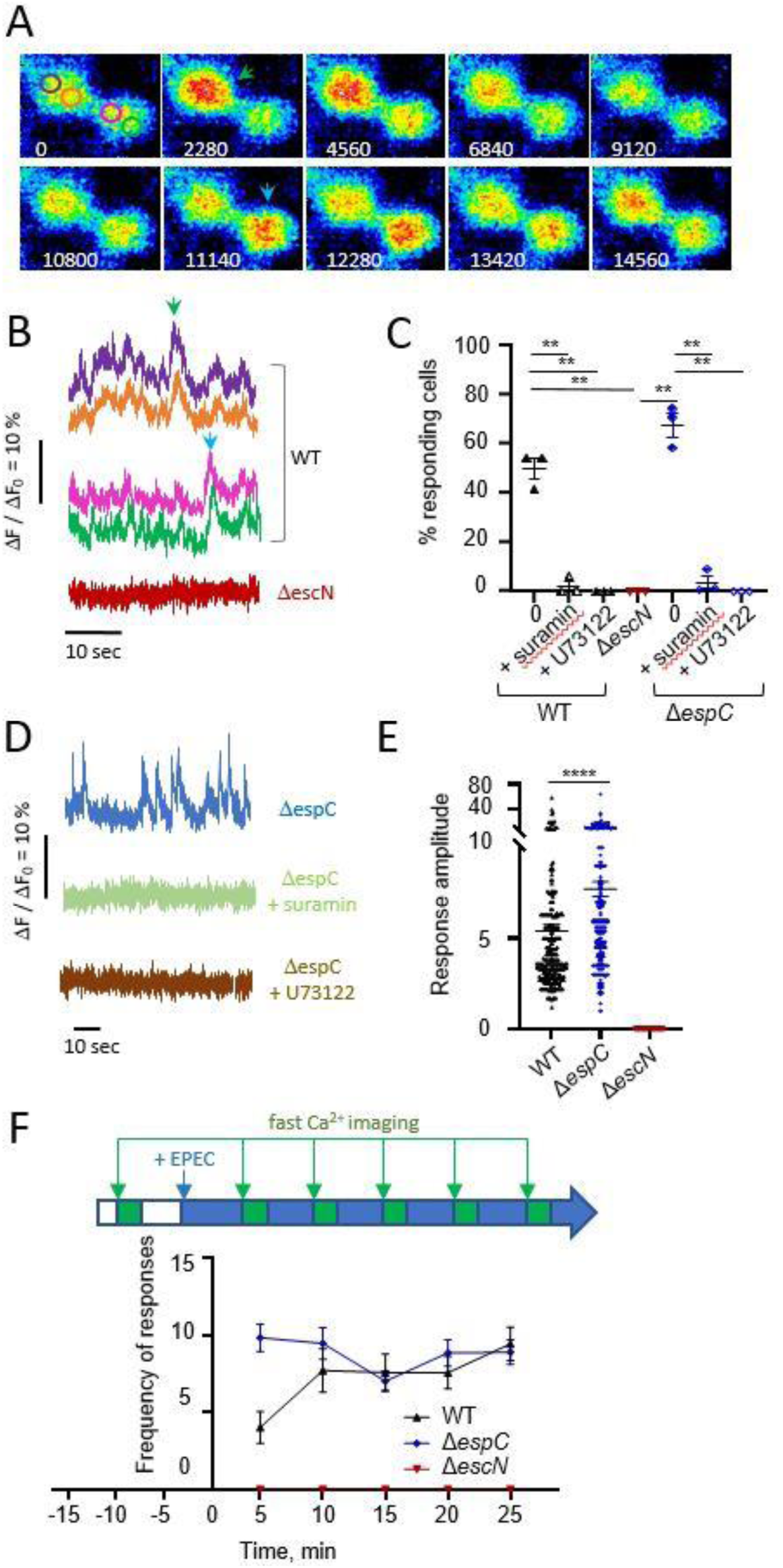
EPEC induces rapid and Coordinated Elementary Ca^2+^ Responses. HeLa cells were loaded with the fluorescent indicator Cal-520, challenged with the indicated bacteria and subjected to high speed Ca^2+^ imaging at a frequency of one acquisition every 57 ms for a duration of 110 seconds. **A**, Representative time-serie of pseudocolored fluorescent micrographs of cells challenged with wild-type EPEC. The numbers indicate the elapsed time in ms from an arbitrarily determined origin. Scale bar = 10 µm. **B**, **D**, traces of Ca^2+^ variations in 2 subcellular regions of the same cell. **B**, WT: traces corresponding to the regions depicted in the image 0 of panel **A**. The arrowheads point to the Ca^2+^ responses shown in Panel A with the corresponding color. **C**, Percent of cells exhibiting Ca^2+^ responses (N > 3, cells > 134). + Suramin: treatment with 200 μM Suramin. +U73122: treatment with 10 μM U73122. **E**, Response amplitude expressed as a percent of the maximal response amplitude induced by treatment with 3 μM histamine (N = 3, cells > 166). **F**, average number of responses per cell. **C**, **E**, Bar: mean. Mann-Whitney test. **: p < 0.01; ****: p < 0.0001. **F**, High speed Ca^2+^ imaging was performed every 5 min for 110 seconds following infection with the indicated bacterial strain as depicted the scheme. The average number of responses per cell is indicated (N > 3, cells > 29).

Together, these results show that at the onset of infection, EPEC induces an atypical pattern of fast Ca^2+^ responses involving the whole or a large area of the cell, likely resulting from low ATP levels released by the insertion of a discrete number of translocons in host cell membranes.

### EPEC-induced fast Ca^2+^ responses are triggered by low ATP levels

Previous studies have described local Ca²⁺ increases triggered by sub-maximal agonist concentrations and leading to limited IP_3_-mediated Ca^2+^ release. These local Ca²⁺ responses are typically small, of short durations and localized to subcellular regions. Among these, the so-called “Blips” correspond to elementary events of opening of a single IP_3_ receptor channel usually lasting between 50 and 100 ms, whereas “Puffs” involve the synchronized activation of multiple IP_3_ receptor channels in localized clusters and last several hundreds of ms (Swillens et al., 1999). In contrast to these described local Ca^2+^ signals, EPEC-induced fast and small responses could occur throughout the cell, suggesting the coordination of elementary responses over large cell area. Since our findings suggested that these responses were elicited by low amounts of eATP released by a discrete number of T3SS translocons, we investigated whether low ATP levels could elicit similar responses.

HeLa cells treated with 150 nM ATP showed Ca^2+^ responses that were indistinguishable from fast responses elicited by EPEC, with an average percent of Ca^2+^ responding cells of 61.2 ± 5.8 % (mean ± SEM) and a frequency of 3.9 responses per cell over 60 seconds (Figs. 4A, C and S4A). These fast Ca^2+^ responses had an amplitude that did not exceed 10 % of the maximal agonist response and occurred over several minutes (Fig. 4A), with a duration of 2.1 ± 1.0 sec (mean ± SEM) (N = 4, 128 responses). As observed for EPEC, fast Ca^2+^ responses induced by low ATP levels involved the whole cell or large cell area encompassing the nuclear and perinuclear area and corresponding to at least 30 % of the cell area as illustrated in Fig. 4B. In this large area, all ROIs corresponding to 1 square micron showed a superimposable profile, as illustrated by traces in Fig. 4C. Similar fast and small coordinated Ca^2+^ responses were also observed when cells were challenged with 100 nM of histamine, another Ca^2+^ agonist, with 44 ± 7 % (mean ± SEM) of cells showing responses, suggesting that these are generic responses triggered by low levels of IP_3_ (N = 2, cells = 340; Fig. S4). EPEC and low concentrations of eATP induced similar responses in polarized intestinal epithelial cells (Fig. S5).

**Fig. 4:**
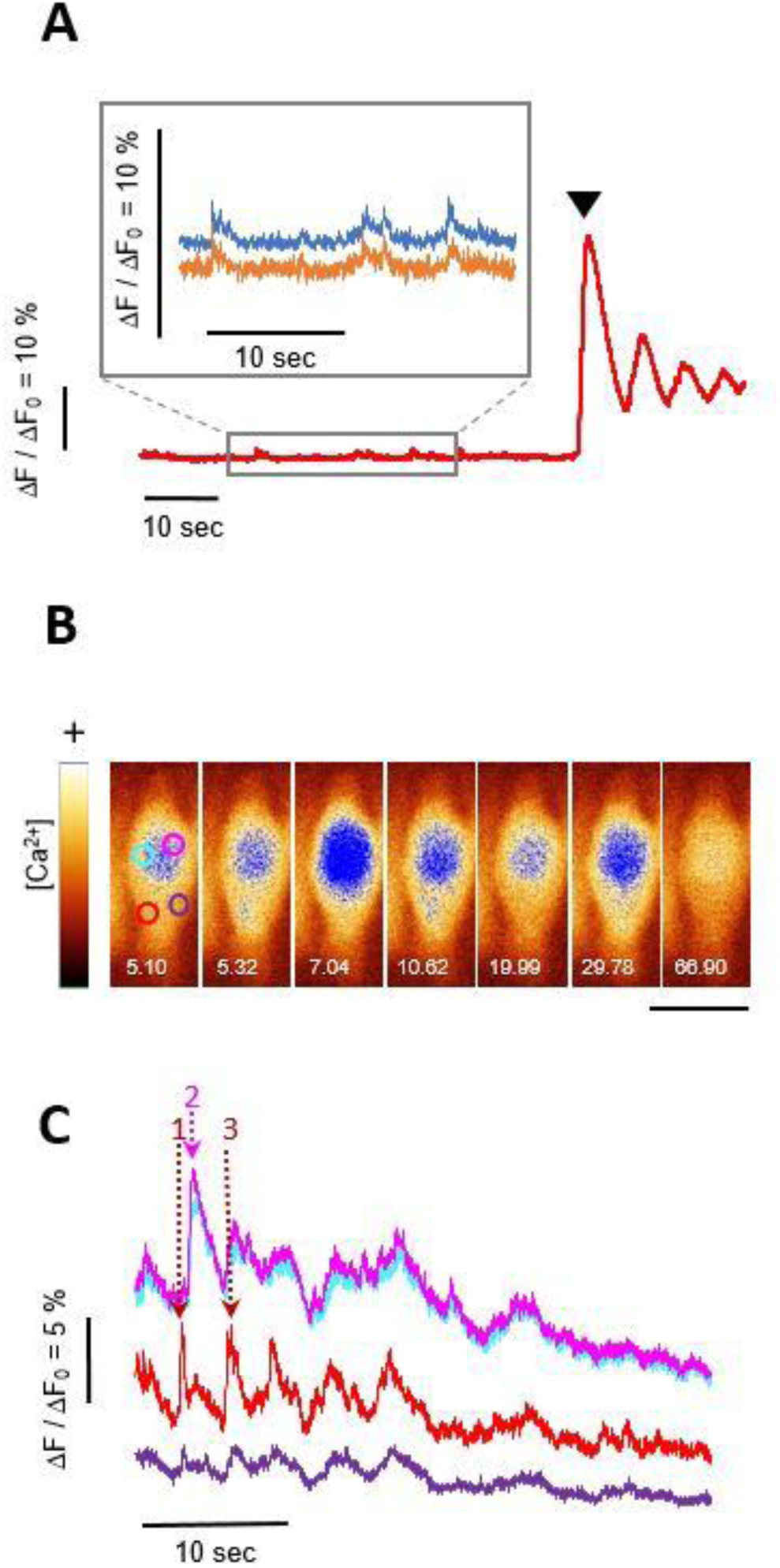
EPEC-induced Coordinated Elementary Ca^2+^ Responses are reproduced by low ATP levels. **A-C**, HeLa cells were loaded with the fluorescent indicator Cal-520, challenged with 150 nM ATP and subjected to Ca^2+^ imaging. Image acquisition every 52 ms (**A**) or 22 ms (**B**, **C**). **A**, Traces of Ca^2+^ variations corresponding to a single cell (red trace), or subcellular regions within the same cell (inset). **A**, arrowhead: challenge with 2 μM ATP. **B**, Time serie of fluorescent micrographs pseudocolored using the “glow” Fiji lookup table, where the blue pixel correspond to an arbitrarily set threshold value. The numbers indicate the elapsed time in seconds. Blue: high intensity pixels showing the large top cell area with CCRICs and the local lower puff area. Scale bar = 10 µm. **C**, Traces corresponding to Ca^2+^ variations in the subcellular regions depicted in Panel **B**. The responses are labelled 1-4, with the response 1 corresponding to the puff (Panel **B**, red ROI) impulsing the response 3 in the same region. Responses 2 and 4 correspond to CCRICs in Panel **B**, blue and green ROIs. Note the diffusion of the responses from the initial release area in other area inferred from the dampening of the response amplitude.

More detailed scrutinizing showed that in their initial mounting phase, these fast Ca^2+^ responses were created by the opening of discrete clusters involving an area of ca. 0.04 μm^2^ that had a transient activity, or possibly were highly mobile, since they were seldom detected at a similar location for three consecutive 22 ms acquisition frames (Figs. S6A, B). These discrete clusters showed similar Ca^2+^ kinetics suggesting the coordination of Ca^2+^ release of single IP_3_R clusters throughout the area that we will hereafter termed CCRICs for “<u>C</u>oordinated <u>C</u>a^2+^ <u>R</u>esponses from IP_3_R <u>C</u>lusters”. Treatment with BAPTA-AM to chelate intracellular Ca^2+^ led to a complete inhibition of CCRICs (N = 3, n > 150 cells; Fig. S7A). Ca^2+^ responses could still be detected upon cell treatment with EGTA-AM consistent with its lower k_on_ rate for Ca^2+^, but with a significant inhibition of the percentage of Ca^2+^ responding cells as well as of the frequency of responses per cell, suggesting that coordination could occur via Ca^2+^ diffusion and Ca^2+^-induced Ca^2+^ release (Fig. S7).

In rare instances (less than 3%), typical local “Puff” responses elicited by these ATP concentrations could also be detected often occurring at the cell periphery (Figs. 4B, red region and 4C, red arrow; Fig. S6D, blue trace) (N > 20, cells > 500). As expected from the small concentrations of Ca^2+^ released at puff sites, no increase in cytosolic Ca^2+^ was detected in a distal cell region (Fig. S6D, top), indicating that isotropic Ca^2+^ diffusion from a puff release site cannot account for Ca^2+^ increase over large cell area. Puffs could also be detected concomitantly with CCRICs in different ROIs of the same cell (Fig. S6D, bottom). In contrast to puffs, CCRICs often showed responses of comparable amplitude in distal regions over the whole cell (Figs. 4C and S6A, B), suggesting the contribution from IP_3_R cluster activation by <u>C</u>a^2+^-Induced <u>C</u>a^2+^ <u>R</u>elease (CICR). Within a given cell, the vast majority of CCRICs appeared quasi-synchronized at the fatest acquisition rate of 22 ms / frame that we could achieve. However, in few instances a delay could be detected in the elicitation of a peak in distant region of a cell (Fig. S6C). These observations suggest that the quasi-synchronization of CCRICs result from the fast diffusion of Ca^2+^ leading to the activation of IP_3_R clusters over large cell area, which may be delayed in a some instances. Scrutinizing of CCRICs showed that while their profiles were comparable, the amplitude of these responses varied in different regions of the cell, with often a single 1 μm^2^ region, likely corresponding the initial firing cluster, showing a prominent amplitude and other regions with smaller amplitude for a given response (Figs. 4B and 4C). For example, in Fig. 4C, the highest amplitude is observed in the red region for peaks 1 and 3, whereas it is observed and in the purple region for peak 2. Thus, for a given CCRIC, the respective contribution of local IP_3_R cluster activation and isotropic diffusion of Ca^2+^from other release sites in Ca^2+^ increase may vary in different regions of the cell.

### CCRICs are coordinated by the rapid diffusion of Ca^2+^ at low concentrations in cell area with a high density of IP_3_ clusters

In Fig. 5, we used modeling to further investigate the mechanism of coordination of these fast Ca^2+^ responses. Based on our previous studies (Voorsluijs et al., 2019; Ornelas-Guevara et al., 2023), the model provides a fully stochastic spatial description of Ca^2+^ release dynamics from IP_3_R clusters in a two-dimensional representation of a HeLa cell. The simulation domain extends on 10 × 10 µm^2^ and is discretized into a 20 × 20 grid of compartments (0.5 × 0.5 µm^2^ each), each representing a cytosolic subvolume of 10^-16^ L. Each compartment contains at most one cluster of IP_3_R, whose dynamics is described as a whole. Besides, cytosolic Ca^2+^ concentration can also vary because of uptake by SERCA, release by a leak or diffusion (Supplementary Information, Note 1).

**Fig. 5.**
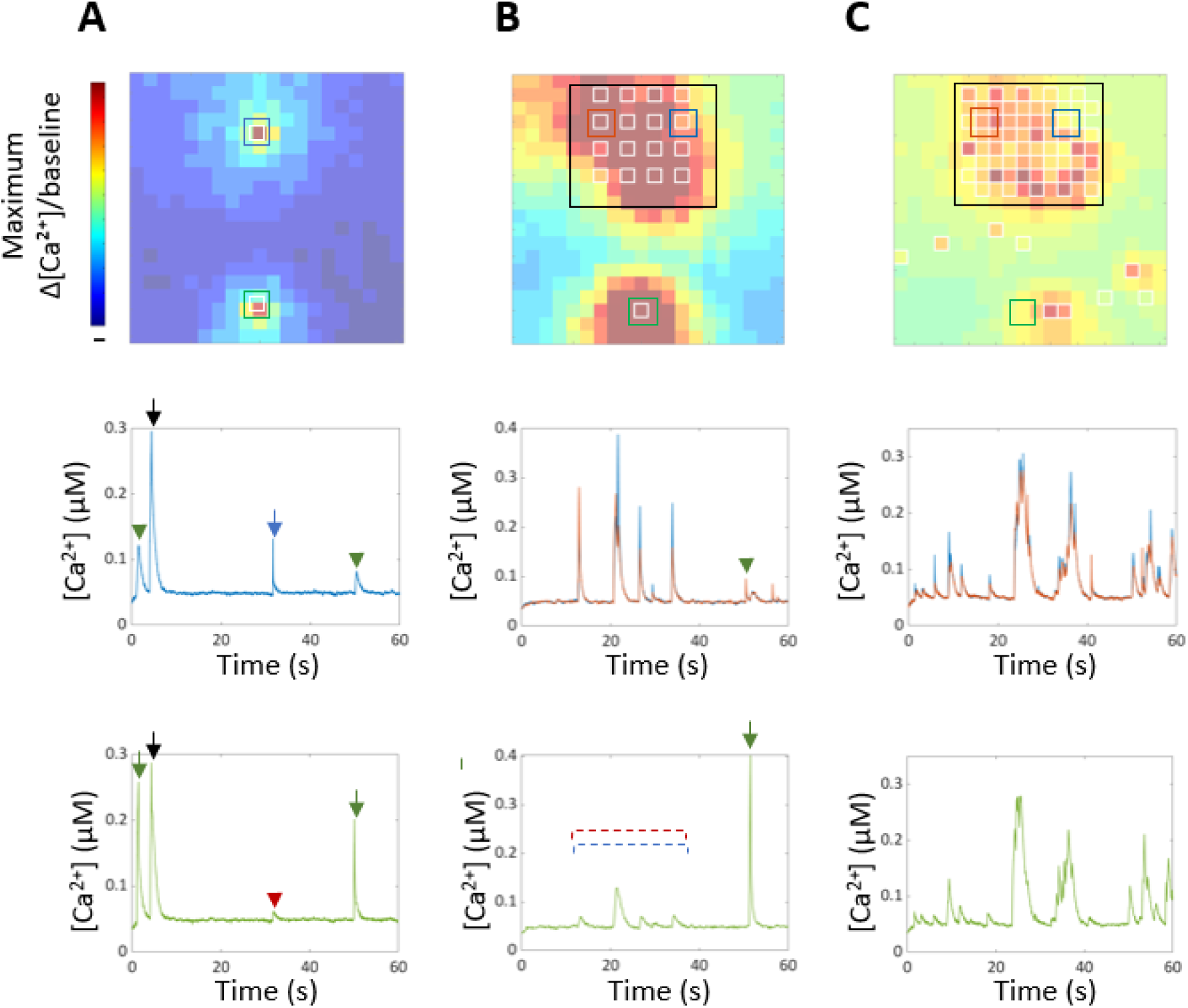
Modeling of Coordinated Elementary Ca^2+^ Responses. **Top,** Ca^2+^ variations in subcellular area within a single cell are represented in pseudocolor. Shown are the maximum values of Δ[Ca^2+^]/[Ca^2+^]_b_ reached in each compartment during a 60s simulation. Empty white squares: IP_3_R clusters. **Graphs**, Traces correspond to Ca^2+^ variations in the region with the matching color. Colored arrows: Ca^2+^ response due to the activation of an IP_3_ cluster in the region with the matching color. Colored arrowhead and dashed red and blue lanes: Ca^2+^ variations due to the diffusion of a Ca^2+^ response from or nearby to the region with the matching color. Black arrows: Ca^2+^ response due Ca^2+^-activated Ca^2+^ release. **A**, low density of IP_3_R clusters with local responses detected. **B**, **C**, Empty black box: area with a high density IP_3_R clusters. **C**, similar to **B**, but following IP_3_R cluster sensitization due to increased Ca^2+^ responses.

In Fig. 5A, low ATP levels lead to low IP_3_ levels activating a limited number of IP_3_ clusters, opening stochastically and releasing small amounts of Ca^2+^. In a given area of the cell, the Ca^2+^ variation integrates Ca^2+^ release from clusters within this area, as well as Ca^2+^ diffusing from or to other cell area. For a given response, the initially firing cluster is contained in an area characterized by the highest Ca^2+^ peak amplitude (Fig. 5A, blue and green arrows) that dampens in a distal area (Fig. 5A, blue and green arrowheads). If the density of IP_3_R clusters is low, as expected for ER compartment at the cell periphery, the spatial segregation of the initial firing cluster and resulting Ca^2+^ diffusion to other area is clearly detected (Fig. 5A). In instances, however, the model predicts temporally coordinated responses of similar amplitude, suggesting Ca^2+^-induced Ca^2+^ release from secondary clusters (Fig. 5A, black arrow). For both type of Ca^2+^ dynamics, a large value of the Ca^2+^ diffusion coefficient of 100 μm^2^ / s is a key parameter that needs to be taken into account in the model. While it is generally admitted that Ca^2+^ diffuses very slowly due to the Ca^2+^ buffers in the cell (∼30 μm^2^/s), the low levels of Ca^2+^ released by CCRICs may not be subjected to the diffusion limitations observed at higher Ca^2+^ levels, because of the relative moderate affinity of buffers for Ca^2+^. How the effective diffusion coefficient of Ca^2+^ is affected by the Ca^2+^ concentration is explained in more detail in Supplementary Information, Note 2.

If the IP_3_R cluster density is high, as expected in the large perinuclear and nuclear area corresponding to the bulk of the ER, the coordination between individual clusters is very fast (Fig. 5B). As a result, the identification of initial firing clusters goes beyond the technical capacities of the imaging set-up, and ROI within this area show comparable profiles of fast Ca^2+^ responses (Fig. 5B). Upon prolonged incubation with increasing Ca^2+^ responses and sensitization of IP_3_R clusters, the coordination of responses linked to Ca^2+^-induced Ca^2+^ release becomes predominant throughout the cell (Fig. 5C). Moreover, at high IP_3_ concentrations, the model reproduces the propagation of a large-amplitude Ca^2+^ wave, as expected (Supplementary Information, Note 3).

### Low eATP levels dampen NF-κB activation

Our findings indicate that the novel Ca²⁺ response pattern is not exclusive to EPEC infection and can be replicated by low levels of eATP or histamine, suggesting a broader physiological relevance. From an immunological perspective, CCRICs may therefore play a critical role in various signaling pathways during bacterial infection. eATP is a well-characterized danger signal contributing to the elicitation of pro-inflammatory signals in various tissues in response to infections (Savio et al., 2018). Previous studies linked intracellular Ca²⁺ signaling and NF-κB activation, a key transcription factor that triggers inflammatory responses (Smedler et al., 2014). We therefore set up to investigate the effects of CCRICs triggered by low eATP levels on NF-κB activation, by performing Western blot analysis against the phosphorylated forms of IκBα (p- IκBα) and P65 (p-P65).

As shown in Figs. 6A and 6B, in control samples, TNF-α induced IκB-α phosphorylation peaking 10 minutes following challenge. In contrast, in the presence of low ATP levels, IκB-α showed a delayed phosphorylation with a 2.1 fold decrease at 10 minutes post-challenge (Figs. 6A and 6B). Consistently, the rates of IκB-α degradation were also slower in the presence of ATP relative to control (Figs. 6A and 6C). As expected from the IκB-α results, TNF-α induced the phosphorylation of the NF-κB P65 subunit and ATP led to a delay and decrease in P65 phosphorylation (Figs. 6D and 6E).

**Fig. 6.**
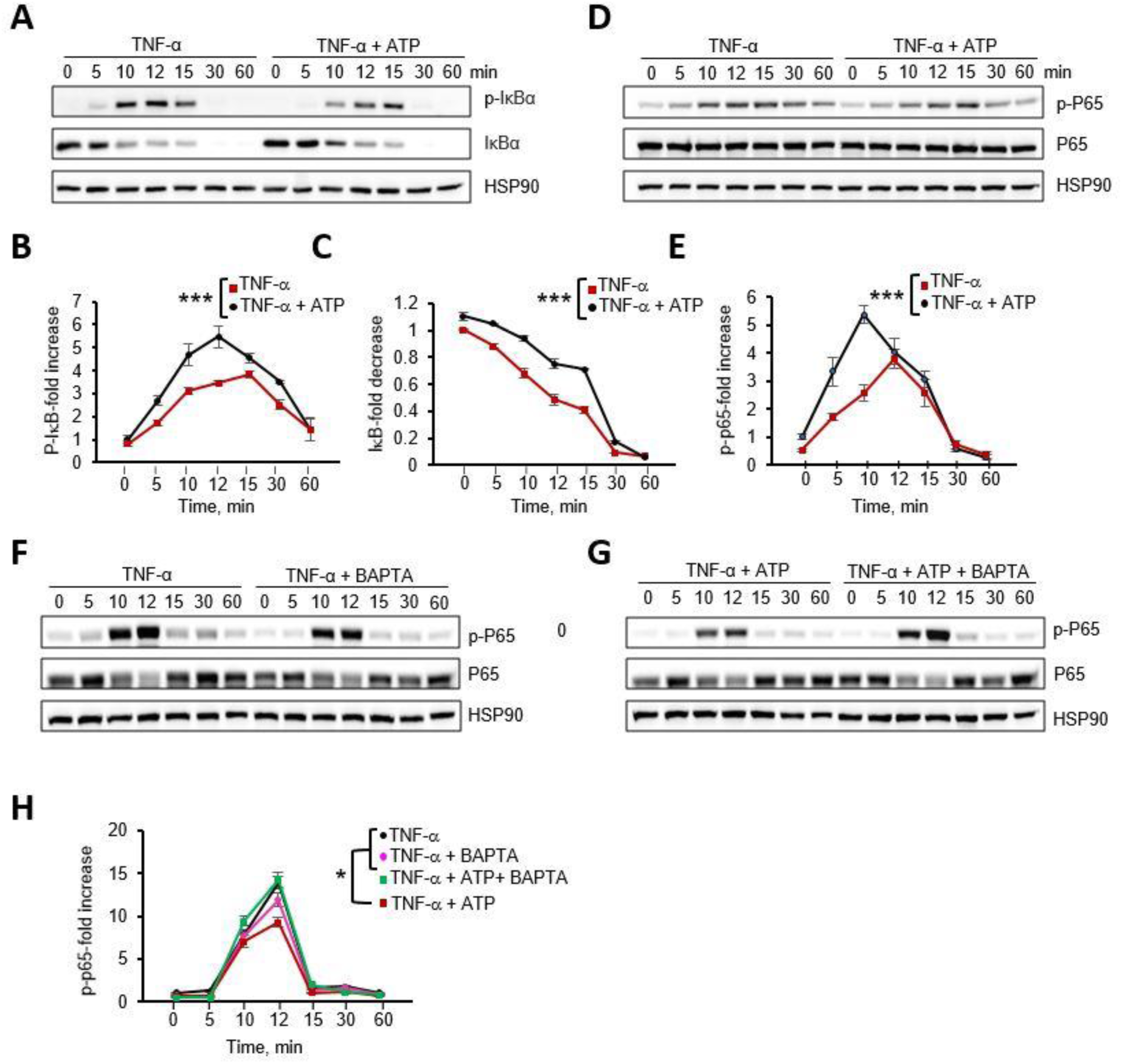
Low ATP levels dampen NF-kappaB activation. HeLa cells were stimulated with 10 ng/ml TNF-α alone or in the presence of 150 nM ATP or 20 μM BAPTA-AM (**F**-**H**). At the indicated time points, cell lysates were analyzed by Western blot using the indicated antibodies**. A**, **D**, **F**, **G**, Representative blots. **B, C, E**, **H**, Densitometry analysis of the indicated antibody signal normalized to that of HSP90 (**B, C**) or total P65 (**E, H**), expressed as fold-increase to basal levels of p-IκB (**B**), IκB (**C**) or p-p65 (**E**, **H**) at time = 0. Values correspond to the mean ± SEM of 3 or 4 independent experiments. p-p65: anti-phospho P65 antibody. p-IκB: anti-phospho IκB antibody. ANCOVA test. *: p < 0.05; **: p < 0.01; ***: p < 0.001.

In control experiments, we did not detect differences in P65 phosphorylation in response to TNF-α stimulation when cells were treated with BAPTA-AM to chelate intracellular Ca^2+^ (Figs. 6F and 6H). However, cell treatment with BAPTA-AM prevented the dampening of P65 phosphorylation triggered by low ATP levels (Figs. 6G and 6H), suggesting that CCRICs down-regulated TNF-α -induced NF-κB activation.

### Low eATP levels down-regulates NF-κB activation through Ca^2+^-dependent O-GlcAcylation

We next investigated how CCRICs could regulate NF-κB activation. O-linked β-N-acetylglucosamine (O-GlcNAc) transferase (OGT) was reported to regulate NF-κB signaling by post-translationally modifying the p65 subunit (Ruan et al., 2017). Interestingly, OGT is regulated by Ca^2+^ signaling, suggesting that CCRICs could affect NF-κB activation via O-GlcNAcylation.

As shown in Fig. S6, TNF-α in the presence of 150 nM eATP stimulated the levels of O-GlcNacylation, specifically for proteins with an apparent molecular weight superior to 100 kDa that was not observed with TNF-α alone. To further investigate the effects of low eATP levels on NF-κB O-GlcNacylation, we performed immunoprecipitation of P65 RelA on lysates of cells stimulated for 12 minutes with TNF-α alone or co-stimulated with TNF-α and 150 nM eATP. As shown in Fig. 7, TNF-α induced increased O-GlcNacylation of P65 relative to non-stimulated cells, but this increase was inhibited by low eATP levels. Inhibition of P65 O-GlcNacylation by eATP was Ca^2+^ dependent, since it was not observed in the presence of BAPTA-AM (Fig. 7).

**Fig. 7.**
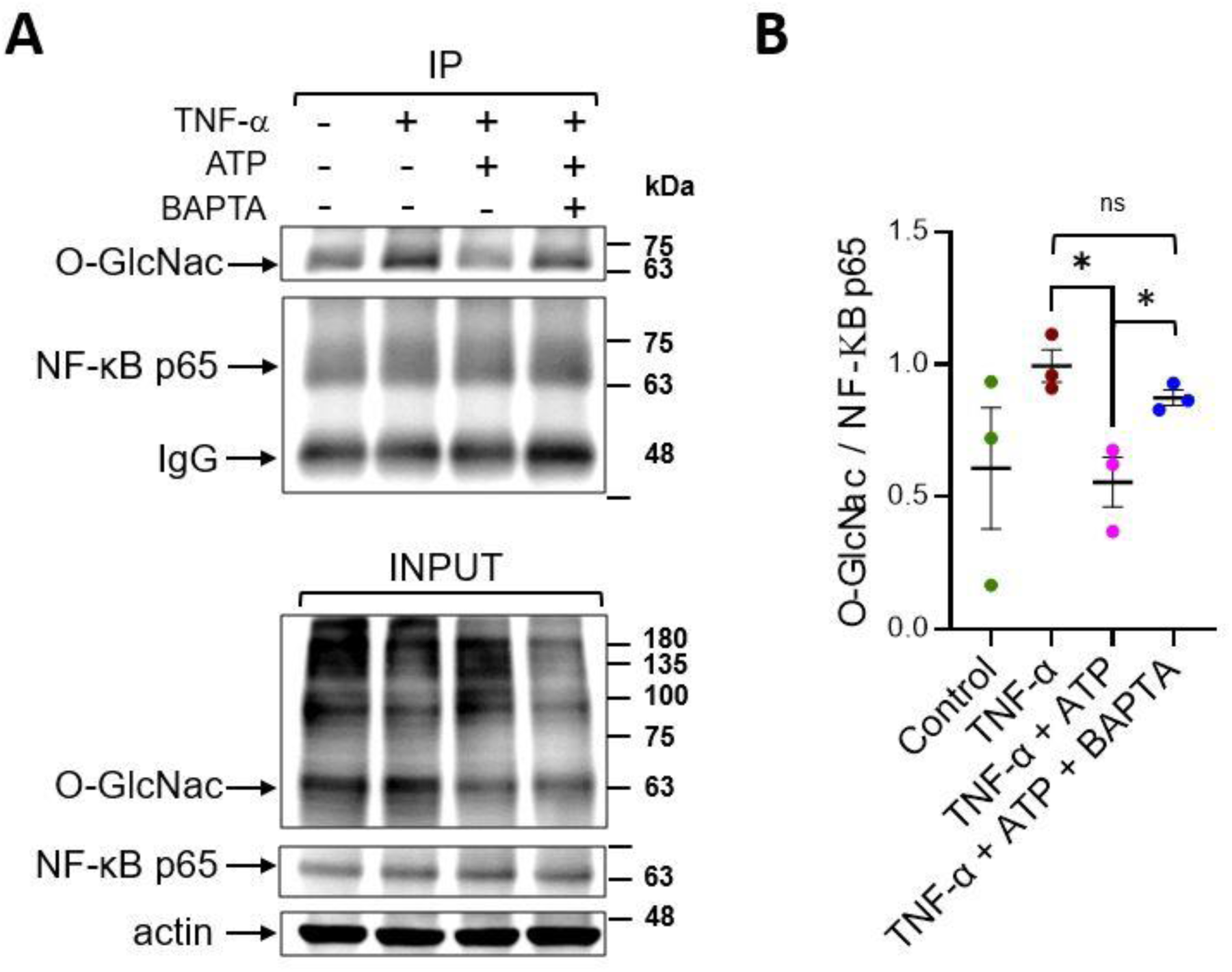
Low ATP levels down-regulate NF-kB O-GlcNAcylation in a Ca^2+^-dependent manner. HeLa cells were stimulated with 10 ng/ml TNF-α alone or in the presence of 150 nM ATP with or without 20 μM BAPTA-AM for 12 min. Cell lysates were subjected to P65 immunoprecipitation. **A**, Representative blots with the indicated antibodies. IP: immunoprecipitates; L: total cell lysates. **B**, Densitometry analysis of the O-GlucNAc signal in P65 immunoprecipitates normalized to that of TNF-α alone. Mann-Whitney test. N = 4. *: p < 0.05. ns: not significant.

The results indicate that low eATP levels inhibit O-GlcNacylation of P65 induced by TNF-α in a Ca^2+^-dependent manner and suggest that differential O-GlcNacylation of P65 relative to higher molecular weight proteins.

## Discussion

We report here the characterization of CCRICs, corresponding to yet undescribed Ca^2+^ responses. CCRICs showed rapid kinetics with an average duration of ca 2.1 seconds and amplitude corresponding to an increase in Ca^2+^ cytosolic concentration of a few hundreds nM, seemingly smaller than that of puffs (Fig. S6D), often occurring repeatedly with a frequency of up to 12 CCRICs / min over the whole cell. Our modelling studies support the notion that CCRICs implicate the rapid coordination of IP_3_R clusters via CICR in large cell area, challenging established concepts in the Ca^2+^ signaling field.

Ca²⁺ diffusion in the cytosol is regulated by mobile and immobile Ca²⁺-binding proteins acting as buffers and forming localized microdomains with steep Ca²⁺ concentration gradients. At the mouth of an open IP₃R channel, Ca²⁺ concentration can reach 100 μM, while just 1–2 μm away, it may drop below 1 μM. Diffusion is restricted by Ca^2+^ buffers with K_D_’s that are generally lower than this concentration. Consequently, Ca²⁺ signaling is generally admitted to be spatially restricted, typically influencing regions within approximately 5 μm of the release site (Foskett et al., 2007). The distribution of Ca²⁺-binding proteins and the spatial arrangement of release channels allow IP₃R-mediated [Ca²⁺]_i_ signals to exhibit diverse spatial and temporal properties, making this system highly adaptable (Vandeput et al., 2007). High-resolution optical imaging of fluorescent Ca²⁺ indicators in intact cells indicates that IP₃-mediated [Ca²⁺]_i_ signals are structured levels (Foskett et al., 2007). At low IP₃ levels, individual IP₃R opens stochastically at discrete release sites, causing localized elevations in cytoplasmic [Ca²⁺]. At higher IP₃ levels, Ca²⁺ release spreads between IP₃R clusters, propagating waves that travel at tens of microns per second, coordinating intracellular signaling ultimately leading to a global Ca²⁺ response. In reference models describing intracellular Ca^2+^ dynamics, cell regions with a high IP_3_R density initiate the Ca^2+^ response from which Ca^2+^ waves propagate with a diffusion coefficient of 10 – 30 μm^2^/s (Falcke et al., 2003).

In contrast to these described global and local Ca²⁺ responses, we found CCRICs to be highly temporally coordinated over large area, suggesting the fast diffusion of Ca^2+^ and propagation of Ca^2+^ by CICR. While challenging generally admitted concepts on the poor diffusion of Ca^2+^, this view is fully supported by our theoretical modeling implicating the fast diffusion of Ca^2+^, with a diffusion coefficient of at least 100 μm^2^/s that can be expected at low cytoplasmic [Ca²⁺]. Indeed, at these low Ca²⁺ concentrations not exceeding a few hundreds nM, the majority of Ca²⁺ buffers are not expected to efficiently bind to Ca²⁺ and to significantly interfere with Ca²⁺ diffusion because of their relative low affinity.

We found that CCRICs implicate a large cell area including the nuclear and perinuclear area. The large CCRIC area likely involves the bulk of the endoplasmic reticulum, while peripheral area may contain smaller ER compartments. In our model considering fast Ca^2+^ diffusion when buffers are far from saturation, the higher density of IP_3_R clusters in the CCRIC area accounts for the high coordination of the responses, relative to the lack of coordination in peripheral area where lower IP_3_R density is expected. Consistently, while CRICCs were detected in the vast majority of cells at these very low agonist concentrations, in rare instances, local “puff-like” responses were also detected at the cell periphery. These observations are in contrast to previously described Ca^2+^ puffs preceding global responses reported to occur preferentially in perinuclear area (Thomas et aL., 1999). These earlier studies, however, involved higher agonist concentrations (1-5 μM ATP) expected to lead to the release of higher IP_3_ concentrations, which may preferentially stimulate larger IP_3_R clusters at the perinuclear region because of the higher density of IP_3_Rs. In addition, larger IP_3_ clusters may release higher amounts of Ca^2+^ for which, as opposed to CCRICs, diffusion would be restrained by Ca^2+^ buffers thereby favoring the spatial confinement of the response.

We showed that a low dose of eATP triggering CCRICs delayed and dampened NF-κB activation linked to a reduction in O-GlcNAcylation of the NF-κB p65 subunit. One major open question is the mechanism by which CCRICS down-regulate NF-κB activation. NF-κB activity can be modulated by O-GlcNAcylation, a reversible glycosylation modification catalyzed by O-GlcNAc transferase (OGT) and O-GlcNAcase (OGA) (Liu & Ramakrishnan, 2021). Previous studies demonstrated that Ca²⁺ signals activate Ca²⁺-regulated enzymes like CaMKII, which in turn phosphorylates and activate OGT, promotes O-GlcNAcylation (Ruan et al., 2017). In other studies, OGT-mediated O-GlcNAcylation could modulate NF-κB signaling pathway (X. Dong et al., 2023). O-GlcNAcylation of P65 could also inhibit its interaction with IκB-α, promote p65 nuclear translocation and increase NF-κB transcriptional activity (Liu & Ramakrishnan, 2021). Reduced O-GlcNAcylation of P65 at residues S550 and S551 was shown to result in decreased NF-κB activation and nuclear translocation (Motolani et al., 2023). These findings are in line with our results suggesting that CCRICs elicited by low-level eATP, downregulate p65 O-GlcNAcylation and NF-κB activation, possibly by modulating OGT activity or OGT-p65 interactions. Reduced O-GlcNAcylation of p65 may affect its phosphorylation patterns indirectly, perhaps by altering the interaction of NF-κB with kinases or phosphatases involved in its activation (Özcan et al., 2010).

Our findings indicate that eATP differentially regulate inflammatory signaling pathways in epithelial cells by dampening NF-κB activation at low levels and stimulating its activation at high concentrations. These results extend the key role of eATP from a danger-associated molecular pattern (DAMP) to a fine-tuner of inflammatory responses depending on its concentration.

## Materials and Methods

### Cell and Bacterial culture

HeLa cells were maintained in Dulbecco’s Modified Eagle Medium (DMEM; Gibco, Thermo Fisher Scientific) supplemented with 10% fetal bovine serum (FBS; Gibco, Thermo Fisher Scientific) Human colon adenocarcinoma Caco-2/TC-7 cells were maintained in DMEM containing 20 % FBS, supplemented with non-essential amino acids. Cells were grown at 37 °C in a humidified incubator with 10% CO₂. Wild-type Enteropathogenic Escherichia coli (EPEC WT), *ΔescN*, and *ΔespC* strains were cultured in Luria-Bertani (LB) broth at 37 °C with kanamycin at a final concentration of 15 μg/ml in a shaking incubator. All strains were transformed with the pmCherry-N1 plasmid, which encodes red fluorescent protein and carries an ampicillin resistance gene for selection. Where applicable, ampicillin (100 μg/mL) and kanamycin were included to maintain resistance markers. The red fluorescence enabled visualization of the bacteria during downstream analyses.

### EPEC infection of cells

HeLa cells were seeded in 6-well plates at a density of 4.5 × 10⁵ cells per well one day before infection. TC-7 cells were seeded at a density of 5 × 10⁵ cells / well in 6-well plates and allow to polarize for 4 days prior to bacterial challenge, replacing medium every day. EPEC WT, *ΔescN,* and *ΔespC* strains grown in the exponential phase were resuspended and primed for 5 hours before challenging HeLa cells in DMEM medium. Cells were challenged with bacteria at an OD600 = 0.2 (Low MOI) or 0.8 (High MOI).

### Ca^2+^ imaging

HeLa cells were seeded onto 25 mm-diameter glass coverslips. Cells were preloaded with the fluorescent indicator dye Cal-520 (AAT #21130) for 30 minutes at room temperature, followed by two PBS washes and one time with DMEM. And placed coverslips, imaging was performed in an observation chamber in DMEM without phenol red, supplemented with 25 mM HEPES. Following a 3-minute baseline acquisition, add the required bacteria strains and OD600 into the chamber. Following a 10-minute incubation at room temperature to allow bacterial attachment. Imaging was then carried out at 35°C to allow for bacterial type III secretion, using a Nikon Eclipse TE200 inverted fluorescence microscope with a 60× objective for 1h of infection. Fluorescence signals were acquired 485 nm with excitation and 535 nm emission parameters. Image control and data acquisition were managed by Simple32 software (Compix Inc.), depending on the experiment requirement, imaging at 22ms, 57ms or 10 sec acquisition intervals. 3 μM of Histamine and 2 μM of ionomycin were applied to check the ability of cells to show calcium response. Images were captured using a CMOS camera (Hamamatsu) and analyzed using the same software.

### Immunofluorescence analysis

Cells were washed three times with PBS and fixed with 3.7% paraformaldehyde (PFA), permeabilized with 0.1% Triton X-100 for 5 minutes and washed with PBS. Blocking was performed using 3% FBS in PBS for 30 minutes at room temperature. Cells were incubated with anti-Z0-1 polyclonal antibody (ThermoFisher Scientific, # 40-2200) for 1 hour at a 1:50 dilution in PBS containing 1% FBS for an hour, followed by anti-rabbit IgG-Alexa 488 (Life Technologies, #A11034) or mouse phalloidin-Alexa 488 (Fischer Scientific, #17511176) at a 1:200 dilution and DAPI (1 μg/ml; Merck, #102362276001) for another hour. Samples were mounted using Dako mounting medium (Agilent) and imaged using a Nikon Ti2 confocal microscope equipped with a 60× objective and Nikon acquisition software.

### Modelling

We develop a fully stochastic spatial model to simulate Ca^2+^ release dynamics from IP_3_R clusters in a two-dimensional representation of a HeLa cell. The simulation domain measures 10 × 10 µm^2^ and is discretized into a 20 × 20 grid of compartments (0.5 × 0.5 µm^2^ each), each representing a cytosolic subvolume of 10^-16^ L. Ca^2+^ exchange between the Endoplasmic Reticulum (ER) and the cytosol occurs through IP_3_R-mediated release, SERCA uptake, and ER Ca^2+^ leak, following the framework by Voorsluijs et al., 2019 and Ornelas-Guevara et al., 2023. Ca^2+^ diffusion is implemented stochastically as a kinetic process between adjacent compartments (Kraus et al., 1996), with a diffusion coefficient of 100 µm^2^/s to reflect moderate endogenous buffering. Each IP_3_R cluster functions as a single unit with four possible states: Open (O), Closed (C) and two Inhibited states (i1, and i2), transitioning in response to local [Ca^2+^] and [IP_3_]. This phenomenological description captures key characteristics of Ca^2+^ puffs and their transition to global signals via Ca^2+^ diffusion and Calcium Induced Calcium Release.

We perform all simulations using the Gillespie algorithm, where each event, reaction or diffusion, is selected stochastically based on its propensity. See Supplementary Information S1 for additional information about the model.

### Western Blot analysis

To obtain total cell extracts, cells were lysed in sample buffer 1x (62.5 mM Tris pH=8, 2% SDS, 10% glycerol, 0.05% bromophenol blue, 5% β- mercaptoethanol) and boiled at 95°C for 5 minutes. Proteins from total lysates were separated by SDS PAGE and transferred to nitrocellulose membrane (0.45μM AmershamTM ProtranTM). Western Blot analysis was performed according to standard procedure using the following primary antibodies diluted in PBS containing 0.1 % Tween-20 and 5 % non-fat milk: IκB-alpha (OZYME, #9424S) at a 1:1000 dilution, Phospho-IκB-α (OZYME, #9424S) at a 1:1000 dilution, NF-κaB p65 (OZYME, #9246S) at a 1:5000 dilution, Phospho-NF-κB p65 (OZYME, #3033S) at a 1:1000 dilution, HSP90 (Santa Cruz Biotechnologies #sc-13119) at a 1:1000 dilution. The secondary HRP-conjugated anti-mouse (Cytiva) and anti-rabbit (Sigma) antibodies were used at a 10-^4^ dilution.

### Immunoprecipitation assays

HeLa cells were seeded in 150 cm^2^ dishes (7.4 × 10⁶ cells/dish) and cultured in DMEM supplemented with 10% FBS. Cells were pretreated with 20 µM BAPTA for 30 minutes at room temperature where indicated, then stimulated with 10 ng/mL TNF-α alone or in combination with 150 nM ATP for an additional 12 minutes. After treatment, cells were washed with ice-cold PBS containing 1 mM NaF and 1 mM Na₃VO₄, lysed in ice-cold lysis buffer (50 mM Tris-HCl pH 7.5, 0.5% Triton X-100, 100 mM NaCl, 1 mM DTT, protease inhibitor cocktail without EDTA), and incubated on a rotating wheel at 4°C for 1 hour. Lysates were clarified by centrifugation at 13,000 rpm for 30 minutes at 4°C. Supernatants were incubated with 5 µL of anti-NF-kB p65 antibody (Abcam, ab16502) for 2 hours at 4°C with rotation, followed by overnight incubation with pre-equilibrated protein A/G beads. Immunocomplexes were washed once with lysis buffer and twice with PBS, then eluted in Laemmli buffer by boiling at 100°C for 5 minutes. Input and IP samples were analyzed by Western blotting using antibodies against O-GlcNAc (Abcam, ab2739), anti-NF-kB p65 (OZYME, 9246S), and Actin (OZYME, 4967).

### Statistical Analysis

All quantitative data are presented as mean ± SEM from at least three independent experiments. Statistical significance was assessed using unpaired two-tailed Student’s t-tests with unequal variance, unless otherwise specified. GraphPad Prism 7 (GraphPad Software) was used for statistical analysis, and p-values < 0.05 were considered statistically significant.

## Acknowledgments

This work was funded by the Inserm, CNRS and ANR grants CalplyCx (ANR-20-CE15-0001), Vital (ANR-24-CE11-3941) to GTVN. FG was funded by a Chinese Science Council PhD fellowship. ROG was supported by Wallonie-Bruxelles International (Excellence Grant 2025). This work was supported by a PDR FRS-FNRS project (T.0073.21). GD is Research Director at the Belgian “Fonds National pour la Recherche Scientifique” (FRS-FNRS).

## Author contribution

FR, LC and GTVN designed, performed experiments and wrote the manuscript. LO performed experiments. ROG and GD performed the mathematical modeling.

## Supplementary Information

### Supplementary Note 1. Description of the computational model

Following previous modeling studies of Ca^2+^ dynamics (Voorsluijs et al., 2019; Ornelas-Guevara et al. 2023), we implement a fully stochastic model of intracellular Ca^2+^ dynamics using the Gillespie algorithm. The model captures IP_3_-induced Ca^2+^ release from IP_3_R clusters, SERCA-mediated reuptake, ER Ca^2+^ leak, and cytosolic Ca^2+^ diffusion. The simulated HeLa cell is represented as a 10 × 10 µm^2^ square, discretized into a 20 × 20 grid. Each compartment (0.5 × 0.5 µm^2^) corresponds to a cytosolic volume of 10^-16^ L.

Each IP_3_R cluster is represented as a single unit with four discrete states: Open (**O**), Closed (**C**), and two inhibited states (**i₁**, **i₂**). The transition from **C** to **O** depend on both [Ca^2+^] and [IP_3_], while the transition from **O** to **i1** depends on [Ca^2+^], following the model described in Ornelas-Guevara et al., 2023 (see Note1. Figure 1). Various spatial distributions of the IP_3_R clusters are investigated in Figure 5A to 5C, where the locations of the clusters are indicated by white boxes. SERCA activity and ER Ca^2+^ leak are present in all compartments.

**Note 1. Figure 1:**
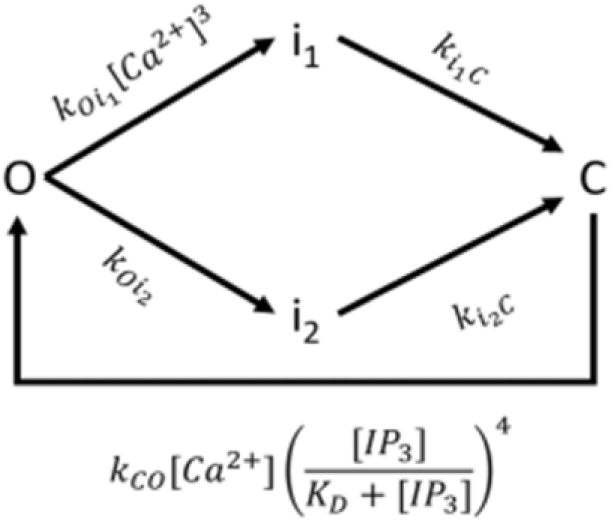
Model describing the dynamics of a cluster of IP_3_Rs. From Ornelas-Guevara et al. (2023). Ca²⁺ diffusion is modelled as a stochastic jump between adjacent compartments (Kraus et al., 1996). The propensity of a Ca²⁺ ion to move depends on the deterministic diffusion coefficient (100 µm²/s) and concentration gradients, assuming isotropic and homogeneous diffusion. All reaction and diffusion events are simulated using the Gillespie algorithm. At each step, an event is selected probabilistically based on its propensity (See propensity functions in Note1. Table 1)

**Note 1. Table 1:**
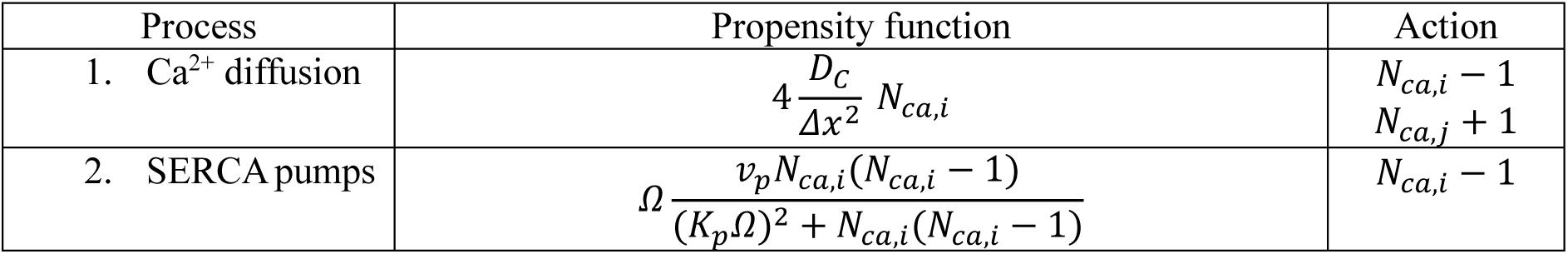
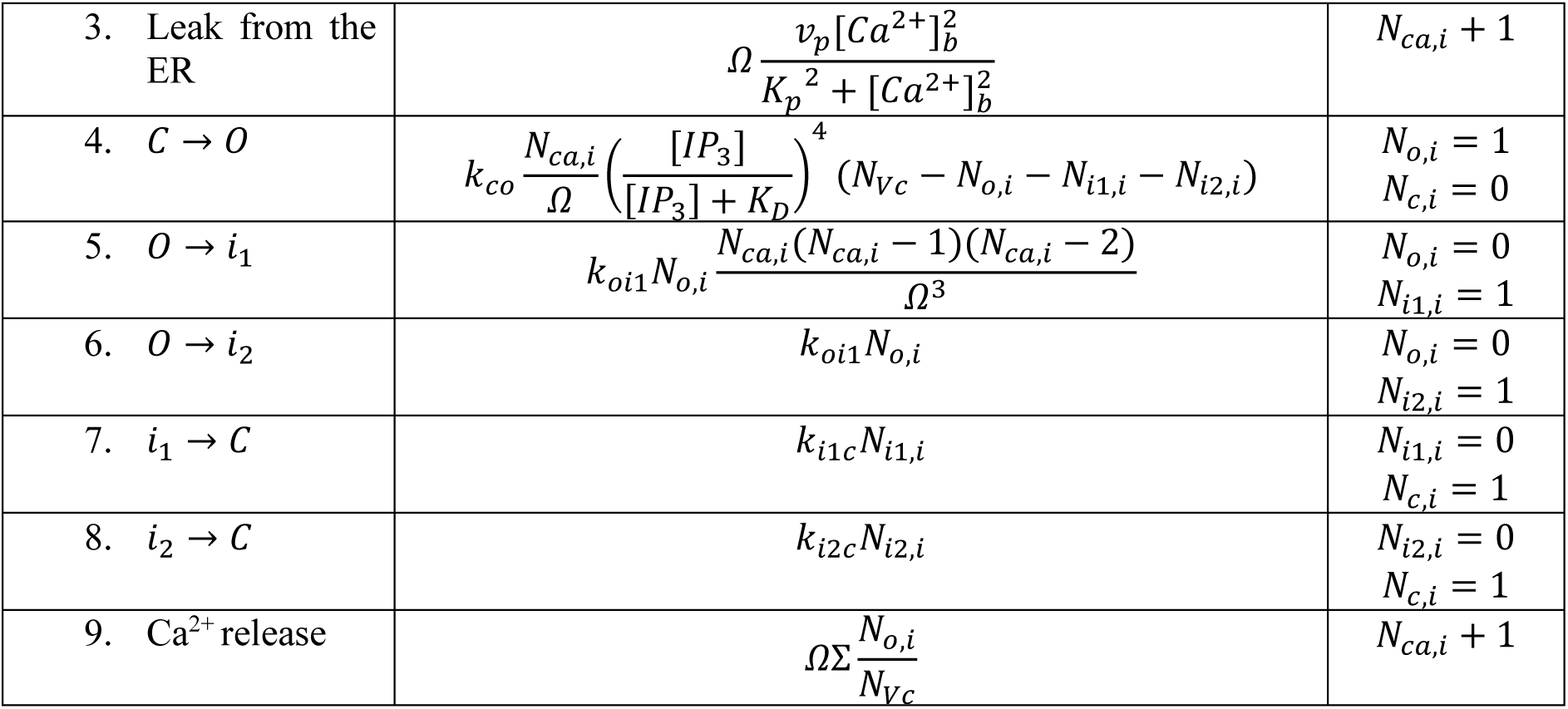
Propensity functions and action of each process. 𝑁_𝑐𝑎,𝑖_ and 𝑁_𝑐𝑎,𝑗_ represent the number of ions in the current box (i) and in an adjacent box (j) selected randomly.

We use an extensivity parameter Ω to convert molecular counts to µM concentrations:

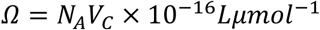

where 𝑁_𝐴_ is Avogadro’s number and 𝑉_𝐶_ is the volume of each compartment.

**Note 1. Table 2:**
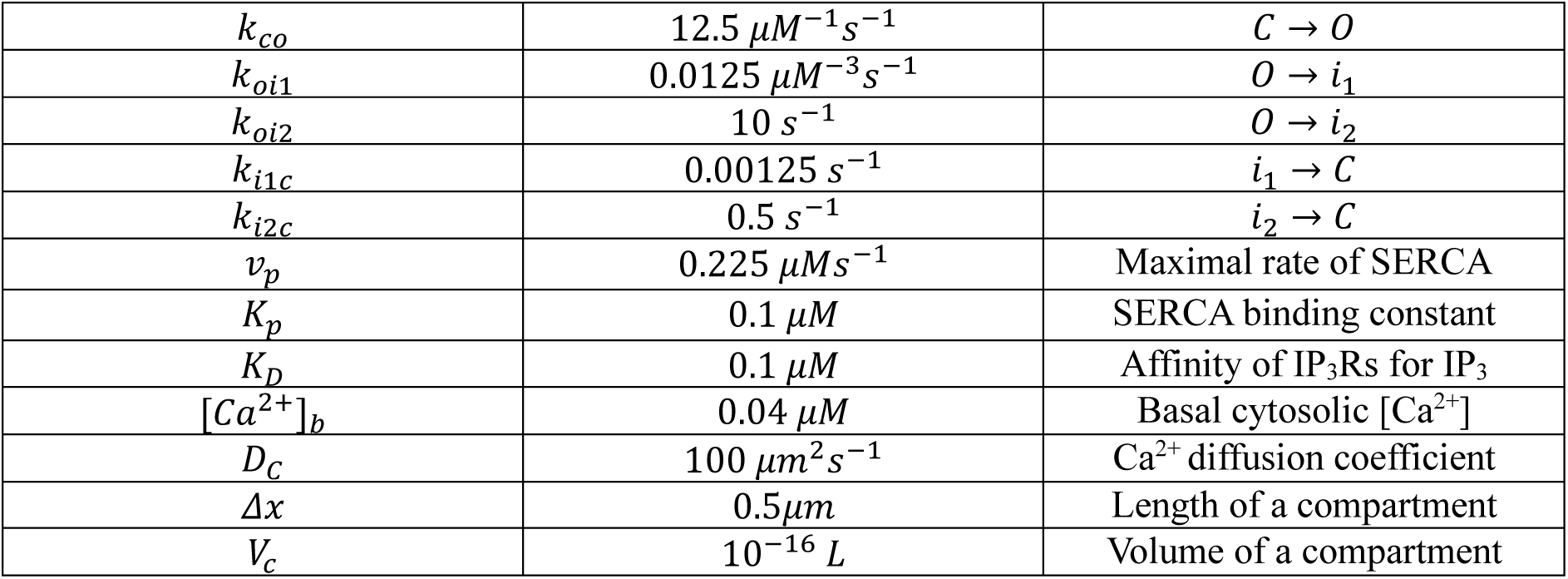
Parameter values used for the simulations shown in Figure 5.

The pseudocolor maps shown in Figure 5 display the maximum Δ[Ca^2+^]/[Ca^2+^]_b_ reached in each compartment during the simulation. These maps are intended to visualize the spatial distribution of the zones of IP₃R-mediated Ca^2+^ release. The algorithm was implemented in MATLAB R2021b.

### Supplementary Note 2. Dependency of the effective Ca^2+^ diffusion coefficient on Ca^2+^ concentration

While in pure water the diffusion coefficient of Ca^2+^ is very large (∼500 μm^2^s^-1^), in the cytoplasm it was reported to be in the range of 13-65 μm^2^s^-1^, mainly due to interactions with Ca^2+^ buffers (Allbritton et al., 1992). This value was determined experimentally by injecting large amounts of ^45^Ca^2+^ at one end of a test tube filled with cytosolic extracts from *Xenopus* oocytes and measuring the spatial spread of radioactivity with time. Such measurements did not assess the dependency of the diffusion coefficient on the Ca^2+^ concentration. A theoretical expression for the effective diffusion coefficient of Ca^2+^ in the presence of buffers under the fast-buffering approximation was derived by Wagner and Keizer (1994). This expression was later shown by Smith et al. (1996) to be valid for a large range of buffering conditions. In this framework, the effective diffusion coefficient is given by

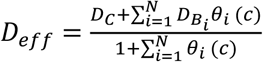

with

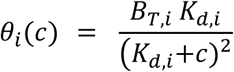

Here, 𝐷_𝐶_ is the diffusion coefficient of free Ca^2+^, 𝐷_𝐵i_ is the diffusion coefficient of buffer species *i*, 𝐵_𝑇,𝑖_ is the total concentration of buffer *i*, 𝐾_𝑑,𝑖_ is its Ca^2+^-binding affinity, and 𝑐 denotes the free Ca^2+^ concentration. The dimensionless quantity 𝜃_𝑖_(𝑐) is the buffering capacity of species *i*.

It is worth mentioning that using the fast-buffer approximation is justified because the Ca²⁺–buffer reaction takes place much faster than Ca²⁺ diffuses over the relevant spatial scales. The characteristic reaction relaxation time can be estimated as τ_relax_ ≈ 1/(k_on_·B_tot_ + k_off_), whereas the diffusion timescale over a distance L is t_D_ ≈ L²/D_c_. For representative values (k_on_ = 100 µM⁻¹·s⁻¹, B_tot_ = 15 µM, K_d_ = 10 µM so k_off_ = K_d_·k_on_ = 1000 s⁻¹), τ_relax_ ≈ 1/(100 µM⁻¹·s⁻¹·15 µM + 1000 s⁻¹) ≈ 4 × 10^-4^ s. Over L = 5 µm, t_D_ = 5^2^/D_c_, which for D_c_ = 220 µm²/s gives t_D_ ≈ 0.11 s. Thus, τ_relax_ is ∼300-fold shorter than t_D_, so for practical purposes the fast-buffer approximation holds in this regime.

Immobile (𝐷_𝐵𝑖i_ = 0) or slowly diffusing buffers (small 𝐷_𝐵𝑖i_) slow down the redistribution of Ca^2+^ and therefore reduce the effective diffusion coefficient 𝐷_eff_ in the Ca^2+^ concentration ranges around and larger than their 𝐾_𝑑_. In contrast, fast buffers (large 𝐷_𝐵𝑖i_) can transport Ca^2+^ away from the source and effectively increase Ca^2+^ mobility. When several buffers with different 𝐾_𝑑,𝑖_, 𝐵_𝑇,𝑖_, and 𝐷_𝐵𝑖_ coexist, their combined effect can make 𝐷_eff_(𝑐) a non-monotonic function of [Ca^2+^], such that effective Ca^2+^ diffusion coefficient can be larger or smaller than the diffusion coefficient estimated in the experiments of Allbritton et al. (1992).

To illustrate how this mechanism can generate concentration-dependent Ca^2+^ mobility, we consider a simple but physiologically plausible mixture of three buffers with distinct properties chosen to be broadly consistent with well-known cytosolic Ca²⁺-binding proteins (e.g., calbindin-D28k, calmodulin, and parvalbumin; Eisner et al., 2023).

· **Buffer 1**: immobile, low-affinity buffer

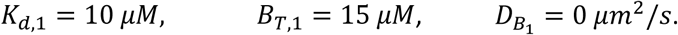

· **Buffer 2**: mobile, moderate-affinity buffer

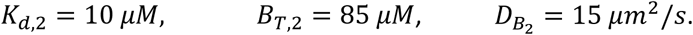

· **Buffer 3**: mobile, high-affinity buffer

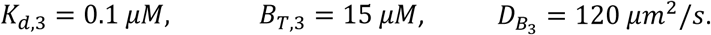

These parameters are not meant to represent specific molecules, but to span a realistic range: an immobile buffer, a generic mobile Ca^2+^ buffer (with a diffusion coefficient on the order of 10–20 µm^2^/s), and a fast-diffusing Ca^2+^ ligand (with a larger diffusion coefficient and lower concentration, conceptually similar to ADP). Many endogenous Ca^2+^-binding species, including both proteins and small metabolites, fall within or between these regimes and, collectively, can act either as “sinks” that confine Ca^2+^ or as “carriers” that help it spread.

Note 2. Figure 1 shows 𝐷_eff_(𝑐) computed from the expressions given above for this buffer mixture. Because different buffers have different 𝐾_𝑑_ and 𝐷_B_, the effective diffusion coefficient value varies with 𝑐 in a non-monotonic way: at some [Ca^2+^] ranges, immobile or slow buffers dominate and reduce Ca^2+^ mobility, whereas at other ranges the contribution of fast mobile carriers is more prominent and increases 𝐷_eff_.

To connect this analysis with spatial [Ca^2+^] profiles, we simulated Ca^2+^ release from a point source in a two-dimensional reaction–diffusion system, including explicit Ca^2+^–buffer binding (i.e. not using the equation to calculate 𝐷_eff_ directly, but the underlying reaction–diffusion equations including Ca^2+^-buffers binding and unbinding with the same parameters as in Note2. Fig. 1A). We considered two cases: a small-amplitude Ca^2+^ release (Note2. Fig. 1B), and a large-amplitude Ca^2+^ release (Note2. Fig. 1C), both with identical buffer parameters and diffusion coefficients. Note 2. Figures 2B and 2C show the resulting [Ca^2+^] profiles 10 ms after release.

This example illustrates that, in the presence of multiple buffers with different kinetics and mobilities, low-amplitude Ca²⁺ elevations can spread further than high-amplitude Ca²⁺ signals. The reason is that low [Ca²⁺] transients are handled mainly by fast, mobile buffers, whereas higher [Ca²⁺] transients increasingly engage slowly diffusing buffers, which then dominate and reduce Ca²⁺ mobility, confining the signal.

Importantly, this is not a universal statement about all possible buffer mixtures, but a specific example showing how realistic combinations of mobile and immobile buffers can produce a situation in which lower-amplitude Ca^2+^ signals have a larger spatial extent than higher-amplitude Ca^2+^ signals.

**Note 2. Figure 1.**
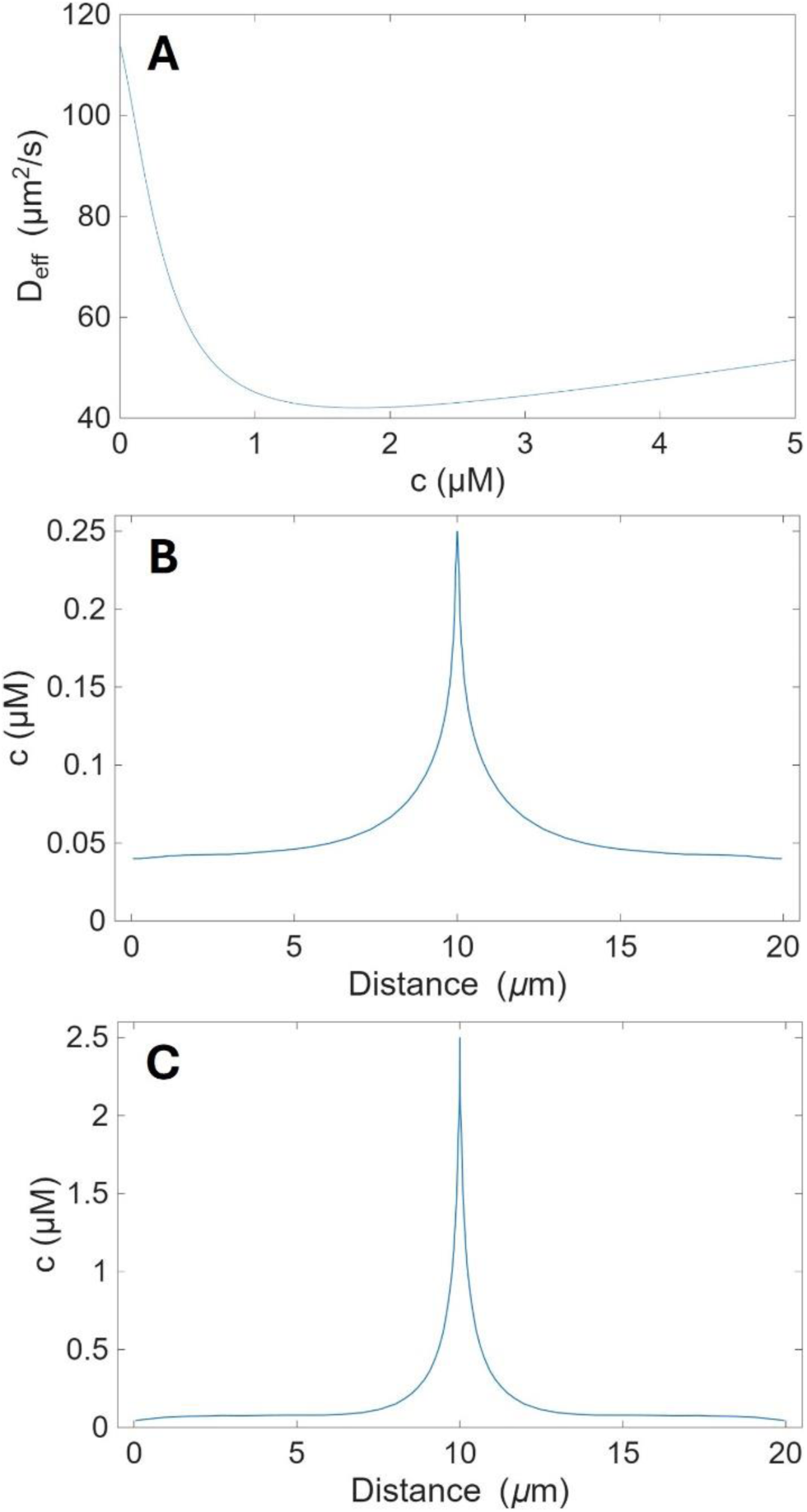
**A**, Effective diffusion coefficient of Ca^2+^ (D_eff_) as a function of free cytosolic [Ca^2+^] (c) for a mixture of three buffers. Buffer 1 is immobile and low-affinity (K_d_ = 10 µM, total concentration 15 µM, diffusion coefficient 0 µm²/s). Buffer 2 is mobile and has moderate affinity (K_d_ = 10 µM, total concentration 85 µM, diffusion coefficient 15 µm²/s). Buffer 3 is highly mobile and has a high affinity (K_d_ = 0.1 µM, total concentration 15 µM, diffusion coefficient 120 µm^2^/s). **B, C,** Spatial profiles of free [Ca^2+^] 10 ms after a constant point-source release in the centre of a 10 × 10 µm^2^ domain representing a cell, obtained from a two-dimensional reaction–diffusion simulation with explicit Ca^2+^–buffer binding using the same buffer parameters as in (A). Only Ca^2+^ release and binding to buffers were included in the simulation. The rate of Ca^2+^ release is 10 times larger in C than in B. [Ca^2+^] peaks at 0.25 µM (**B**) and 2.5 µM (**C**).

### Supplementary Note 3. Global Ca²⁺ responses in the IP₃R cluster model at higher stimulation and stronger Ca^2+^ buffering

In the stochastic cluster model used in Fig. 5, IP_3_R clusters are represented at discrete spatial sites. Each cluster senses the local Ca²⁺ concentration and its stochastic gating depends on this local [Ca²⁺] and on [IP_3_]. Buffers are not included explicitly. Instead, Ca²⁺ diffusion in the cytosol is described by 𝐷_eff_, which accounts for the combined action of endogenous Ca²⁺-binding species.

In the regime relevant for CCRICs, we use 𝐷_eff_ = 100 µm^2^/s and a sub-threshold [IP_3_] = 0.07 µM, below the level that produces global Ca²⁺ responses in the model. With these values the model reproduces key properties of CCRICs: fast kinetics, small amplitude, and large spatial extent.

To simulate the behaviour of the model at higher IP3 stimulation levels, above-threshold IP_3_ concentration, we increased [IP_3_] to 0.1 µM (kept constant in time and space) and used a smaller effective diffusion coefficient, 𝐷_eff_ = 40 µm^2^/s expected for the stronger buffering and lower Ca²⁺ mobility with higher-amplitude Ca²⁺ signals (Note 2. Fig. 1).

Under these conditions, the same cluster model generates a global Ca²⁺ response with larger amplitude and longer duration, rather than a loss of activity due to excessive inhibition of the clusters (Figure S3A). Thus, within a single modelling framework, low [IP_3_] and 𝐷_eff_ = 100 µm^2^/s give rise to CCRIC-like responses, whereas higher [IP_3_] and a more strongly buffered regime (𝐷_eff_ = 40 µm^2^/s) produce robust global Ca²⁺ signals, comparable to global responses observed experimentally (Note 3. Fig. 1).

**Note 3. Fig. 1.**
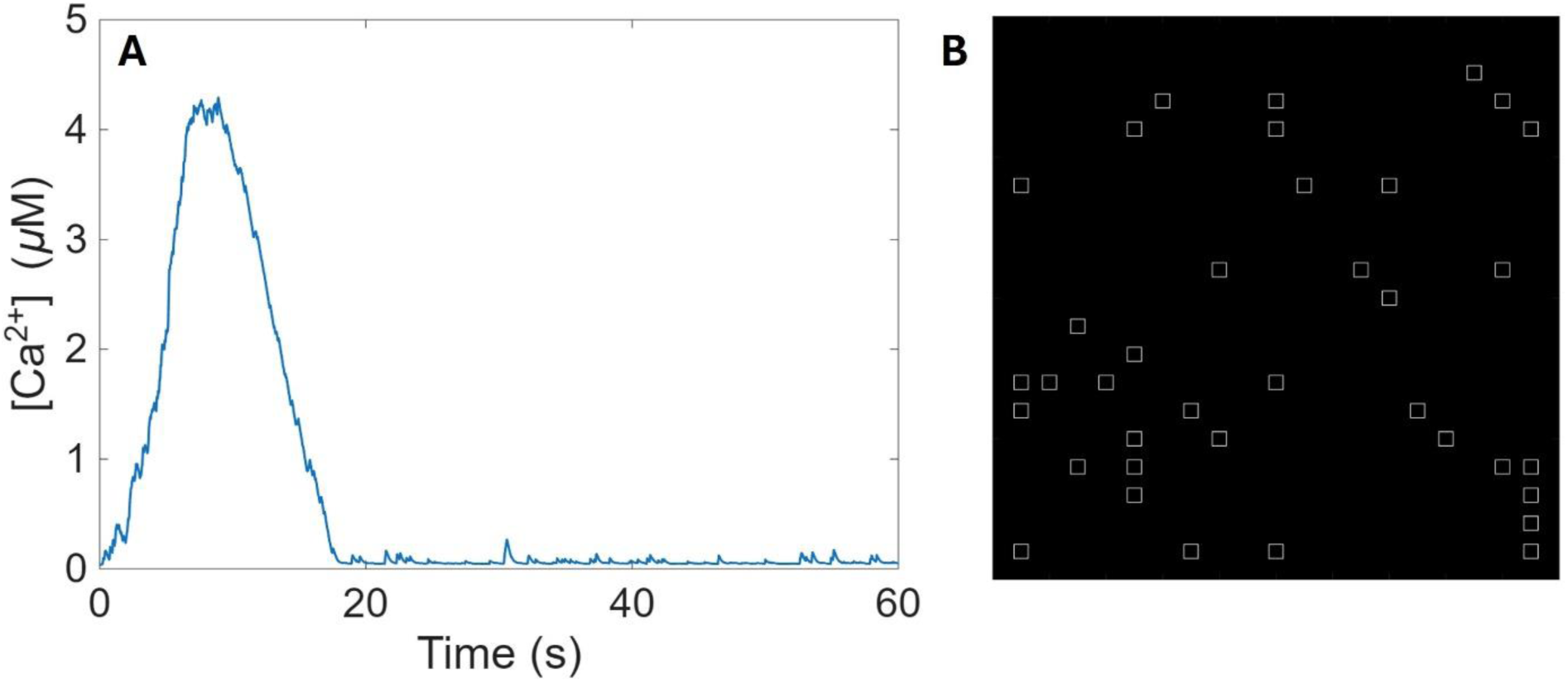
Global Ca²⁺ response at higher [IP_3_]. **A**, Time course of the averaged cytosolic [Ca²⁺] obtained from the stochastic IP_3_R cluster model for a uniform [IP_3_] = 0.1 µM and an effective diffusion coefficient 𝐷_eff_ = 40 µm^2^/s. All other parameters are identical to those used in the simulations shown in Fig. 5. Under these conditions, as opposed to CCRICs, the model produces a “classical” global Ca²⁺ response with larger amplitude and long duration. **B,** 2D cell geometry (10 × 10 µm²) used in the simulations. Squares indicate the random positions of IP₃R clusters.

---

**Fig. S1.**
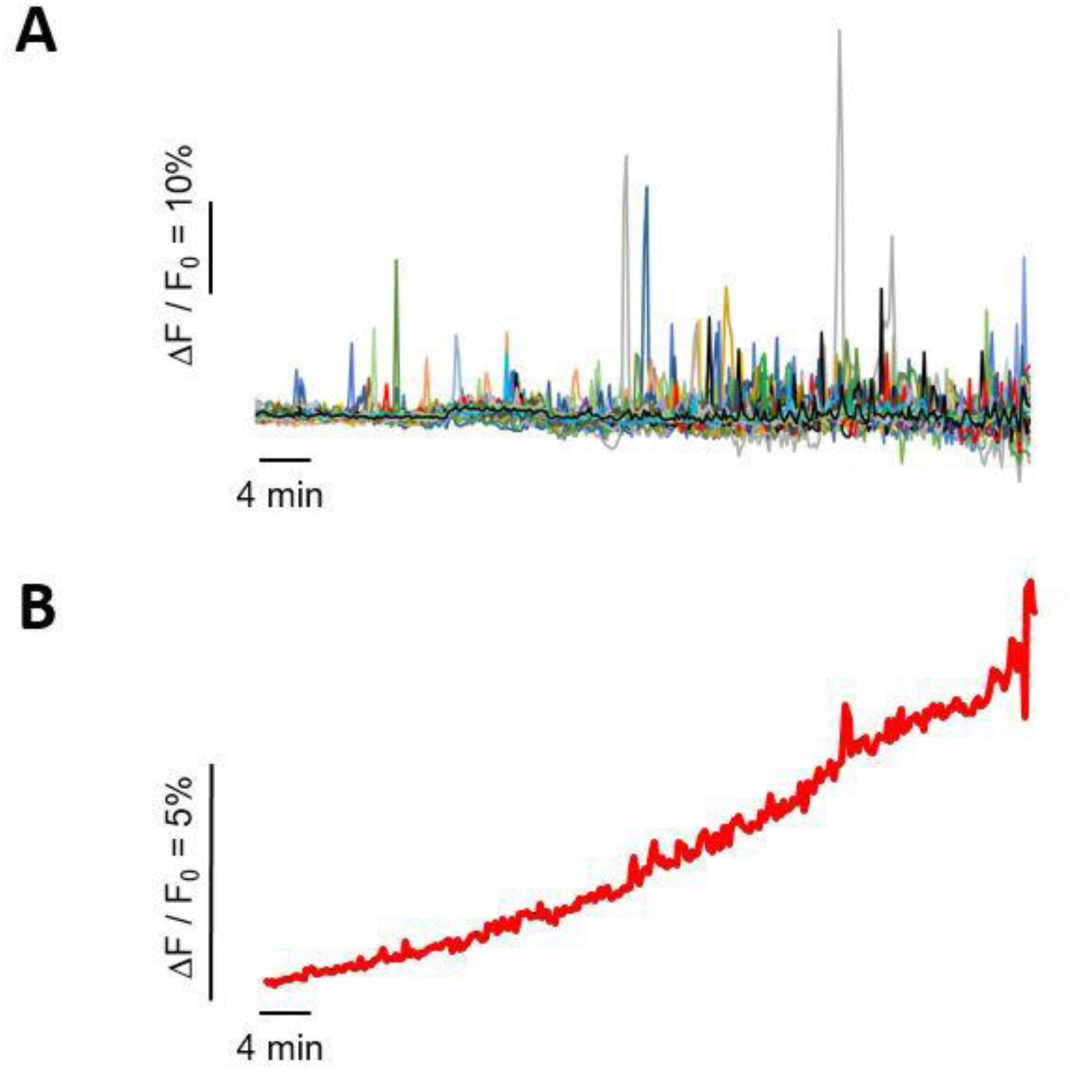
EPEC induces Ca^2+^ responses with a frequency increasing during the time of infection. Ca^2+^ imaging was performed on HeLa cells loaded with the Ca^2+^ fluorescent indicator Cal-520 and challenged with EPEC at a MOI of 100. A, Traces of Ca^2+^ variations in single cells. B, Trace corresponding to the average of traces shown in B (N = 2, cells = 43). Note that cells do not show an increase in basal Ca^2+^ levels that could be mistakenly interpreted from the averaging of Ca^2+^ responses over the cell population.

**Fig. S2.**
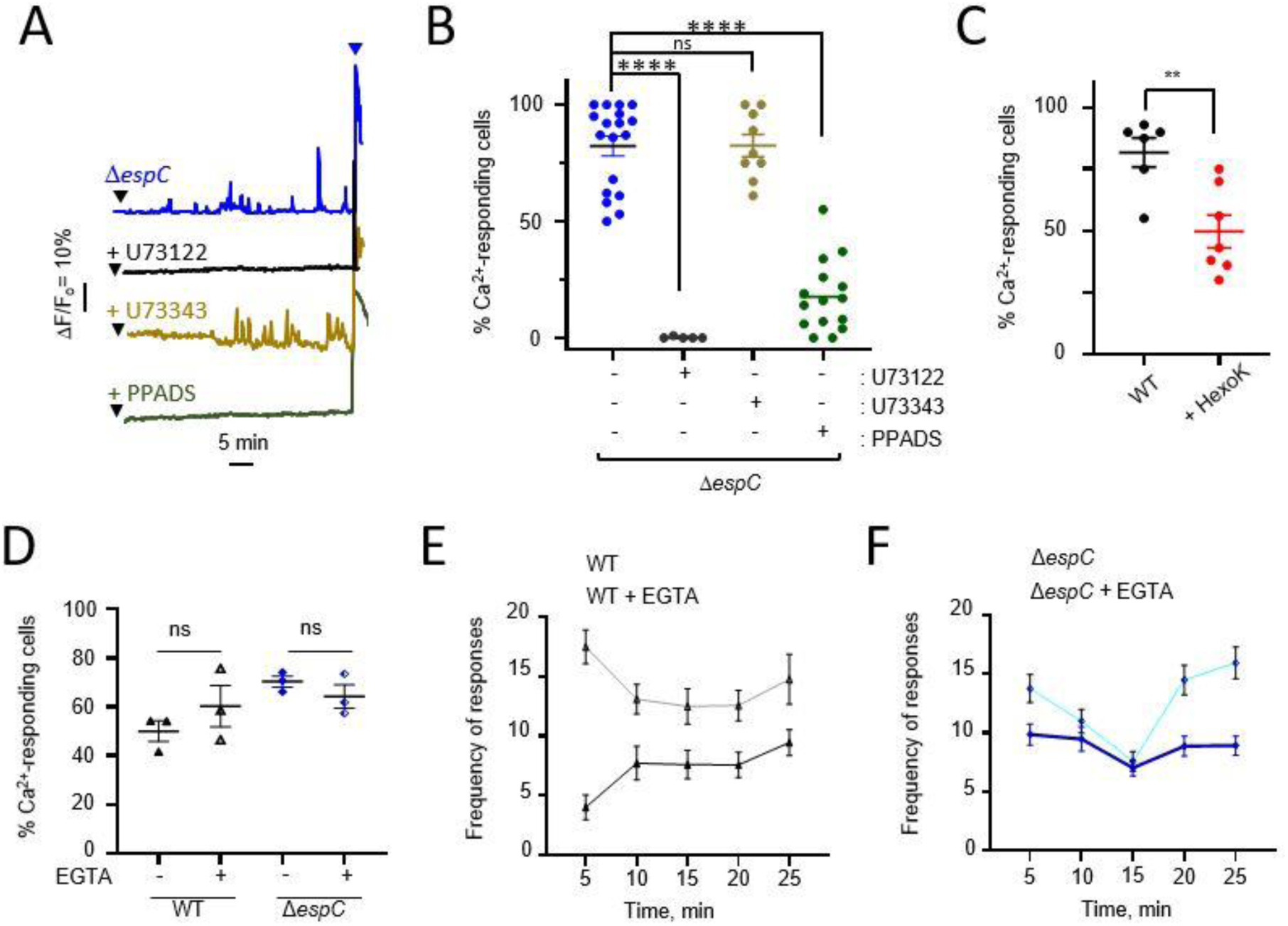
EPEC-induced Ca^2+^ responses does not depend on Ca^2+^ influx but on Ca^2+^ release. HeLa cells were loaded with the fluorescent indicator Cal-520, challenged with a high MOI of EPEC wild-type (WT) or a low MOI of the *ΔespC* mutant and subjected to live-cell Ca^2+^ imaging at a frequency of one acquisition every 10 seconds (**A-C**). **A**, Representative traces of Ca^2+^ variations in single cells. The black arrowheads indicate the time of bacterial challenge. The blue arrowheads indicate stimulation with 3 µM histamine. **B**, Percent of cells exhibiting Ca^2+^ responses. Cells challenged with: *ΔespC* (N = 4, cells > 400); + U73122: *ΔespC* in the presence of 10 μM U73122 (N = 4, cells > 160); + U73343: *ΔespC* in the presence of 10 μM U73343 (N = 2, cells = 230); + PPADS: *ΔespC* in the presence of 20 μM PPADS (N = 4, cells > 400). **C,** cells challenged with wild-type EPEC in the absence or presence of + hexokinase (200 units/ml) and 5 mM glucose (+ HexoK, N = 3, cells > 120). **D-F,** + EGTA: cells treated with 4 mM EGTA. (N = 3, cells > 30). **D**, Percent of cells showing Ca^2+^ responses. **E**, **F**, Frequency of Ca^2+^ responses per cell. Mann-Whitney test. ns: not significant. **: p < 0.01; ****: p < 0.0001.

**Figure S3.**
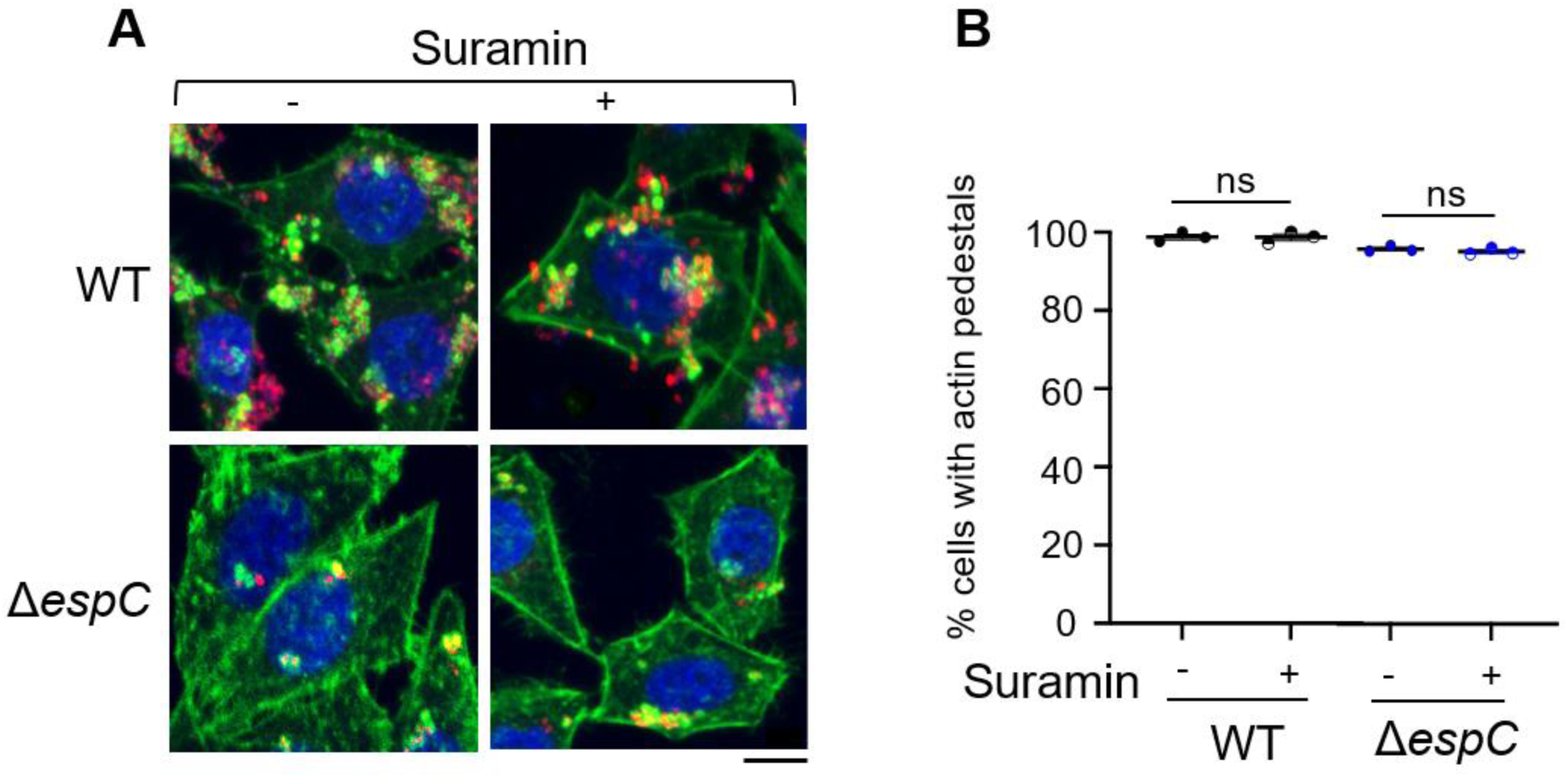
Suramin does not affect EPEC-Induced actin pedestals. HeLa cells were challenged with the indicated RFP-expressing bacterial strains in the presence or absence of 100 μM suramin. Samples were fixed and processed for fluorescence staining. **A**, Representative micrographs. Blue: DAPI; green: Phalloidin-Alexa488; red: bacteria. Scale bar = 10 µm. **B**, Percent of cells exhibiting actin-rich pedestals. High MOI: 50 bacteria / cell. Low MOI *ΔespC*: 10 bacteria / cell. (N=3, n > 150). Mann Whitney test. ns: not significant.

**Fig. S4.**
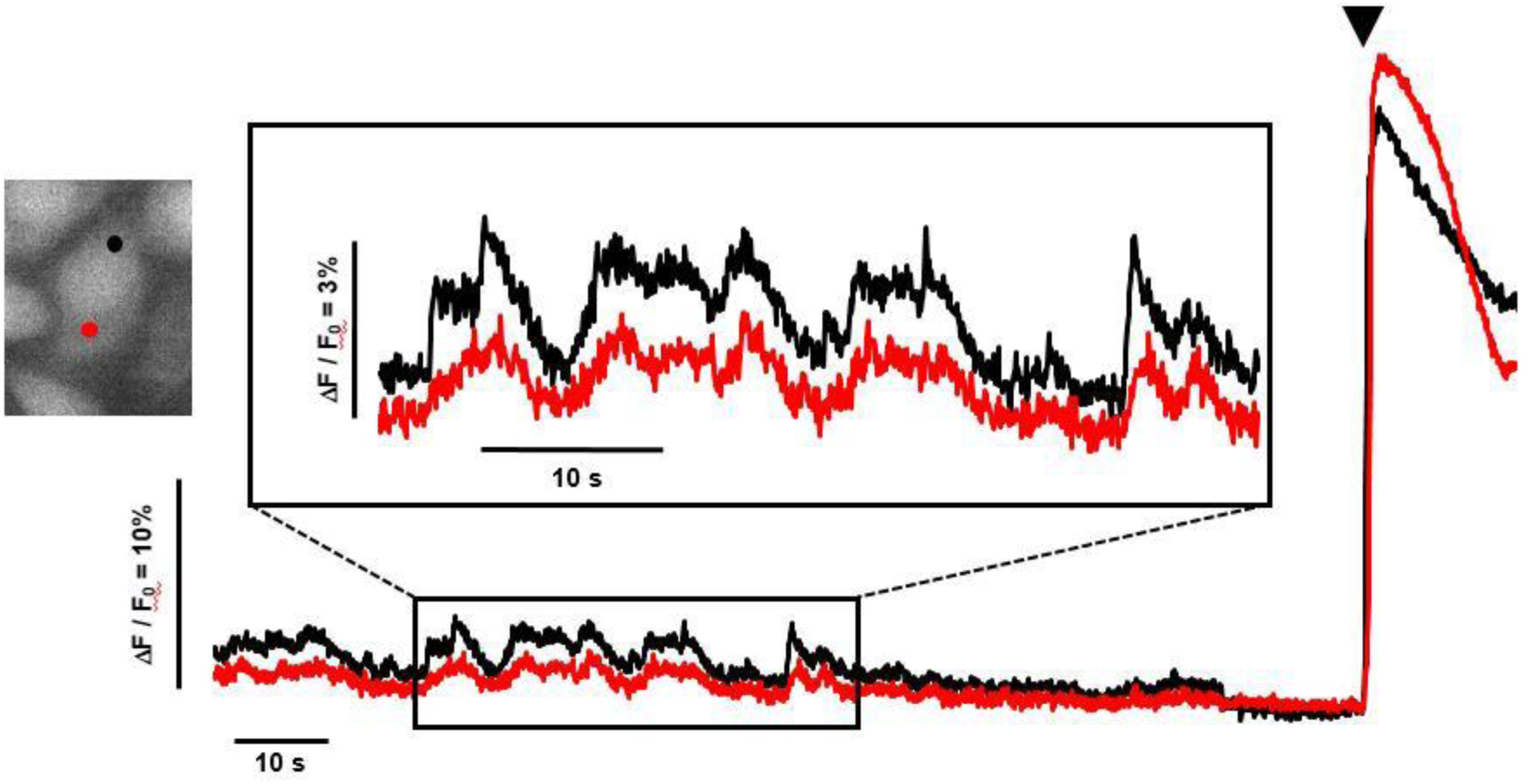
Low concentrations of histamine also induce small and fast coordinated Ca^2+^ responses. HeLa cells were loaded with the fluorescent indicator Cal-520, challenged with 100 nM histamine and subjected to Ca^2+^ imaging, with image acquisition performed every 57 ms. Left: fluorescence micrograph of the single cell with black and red ROIs corresponding to the traces of Ca^2+^ variations in matching color. The box corresponds to a higher magnification of the inset in traces shown at the bottom. Arrowhead: challenge with 10 μM histamine. Traces are representative of 340 cells analyzed from 2 independent experiments, with a frequency of 4.5 ± 0.4 peaks·min⁻¹ (mean ± SEM) and peak duration of 4.45 ± 0.19 s (mean ± SEM).

**Fig. S5.**
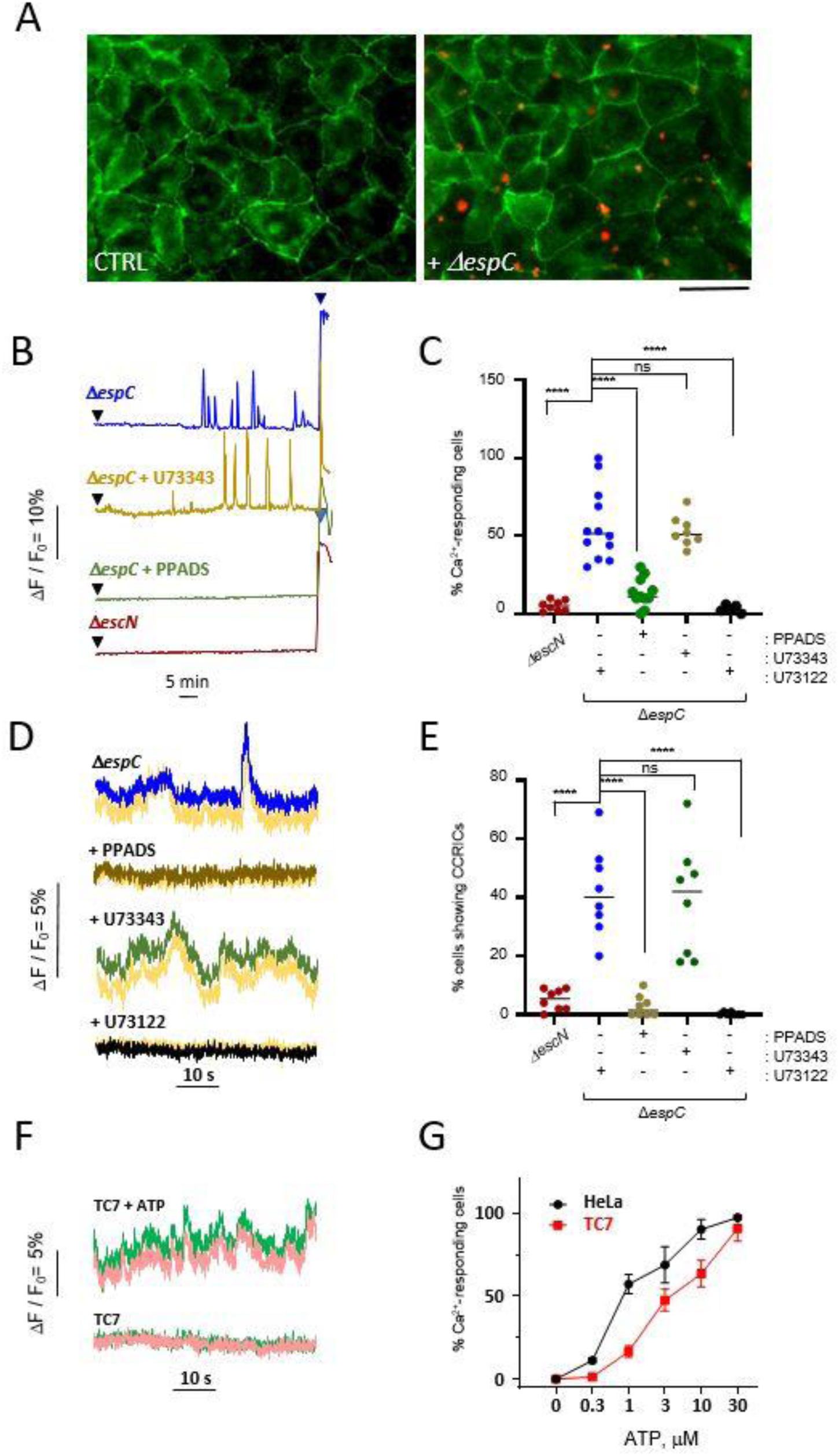
EPEC induces CRICCs in polarized intestinal cells via eATP release. Polarized TC7 cells were loaded with the fluorescent indicator Cal-520 were challenged with the EPEC *ΔespC* strain expressing the Red Fluorescent Protein (RFP) (**A**-**E**) or with ATP (**F**, **G**). **A**, Representative confocal micrographs. Samples were, fixed and processed for fluorescence microscopy analysis. Green: ZO-1 staining. Red: RFP fluorescence. Scale bar = 10 µm. **B-G**, Samples were subjected to live-cell Ca^2+^ imaging at a frequency of one acquisition every 5 seconds. **B**, **D**, Representative traces of Ca^2+^ variations in single cells challenged with the indicated strain and inhibitor. The black arrowheads indicate the time of bacterial challenge. The blue arrowhead indicates stimulation with 3 µM histamine. **C**, **E**, Percent of cells exhibiting Ca^2+^responses (**C**) (N = 4, cells > 241) or CCRICs (**E**) (N =3, cells > 66) following challenge with the indicated bacterial strain and inhibitor. Each value corresponds to a replicate. Mann-Whitney test. ****: p < 0.0001. **F**, Representative traces of Ca^2+^ variations following challenge with 150 nM ATP (TC7 + ATP) or in buffer alone (TC7). The pink and green traces correspond to Ca^2+^ variations in ROIs within the same cell. **G**, Percent of cells exhibiting Ca^2+^responses following challenge with the indicated ATP concentration. Each value represents the mean ± SEM of at least 140 cells in 3 independent experiments.

**Fig. S6.**
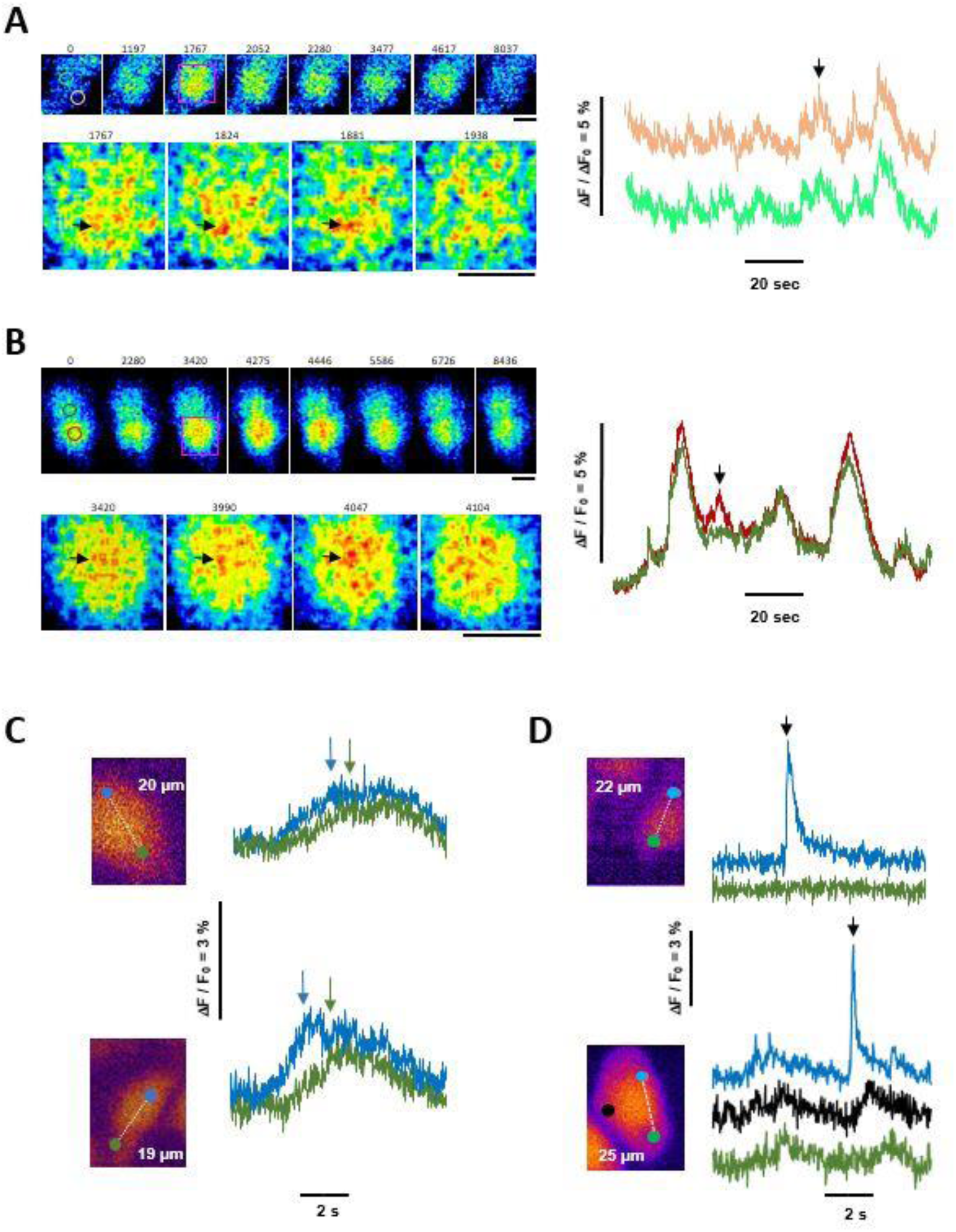
CCRICs induced by EPEC result from Ca^2+^ released by highly transient or mobile IP_3_R clusters. HeLa cells were loaded with Cal-520 and challenged with wild type EPEC at a MOI of 50 bacteria / cell. **A**, **B**, Globally Coordinated responses associated with small and highly mobile clusters. **Time series micrographs**, Cal-520 intensity depicted in pseudocolor. Numbers: time in ms. Lower Panels are magnification of the purple box in the upper panel, with a time interval of 57 ms. Scale bar = 5 μm. Note the high mobility of small clusters / channels. The black arrows point at larger and less mobile clusters. **Traces**, variations of Ca^2+^ in ROI depicted at T = 0 in the micrographs in the corresponding color. The black arrow points at the response peak illustrated by the time series. **C**, **D**, micrographs, Cal-520 fluorescence depicted in the Fiji red lute. Images were taken every 22 ms. Solid circles: ROIs where the variations of Ca^2+^ are shown in traces in the corresponding on the right. The blue ROI correspond to the Ca^2+^ release source based on the higher response amplitude, and the green ROI to a distal region. The number indicates the distance between the ROIs shown by the dashed line. **C**, Arrows point at the peak of the response in the ROI with the corresponding color. Note the delay in response elicitation between the source- (blue) and distal (green) ROI. **D**, Puff-like responses (black arrowhead). Note the absence of Ca^2+^ variations in the distal region (Top), and the simultaneous elicitation of a puff and CCRICs in different ROIs (Bottom).

**Fig. S7.**
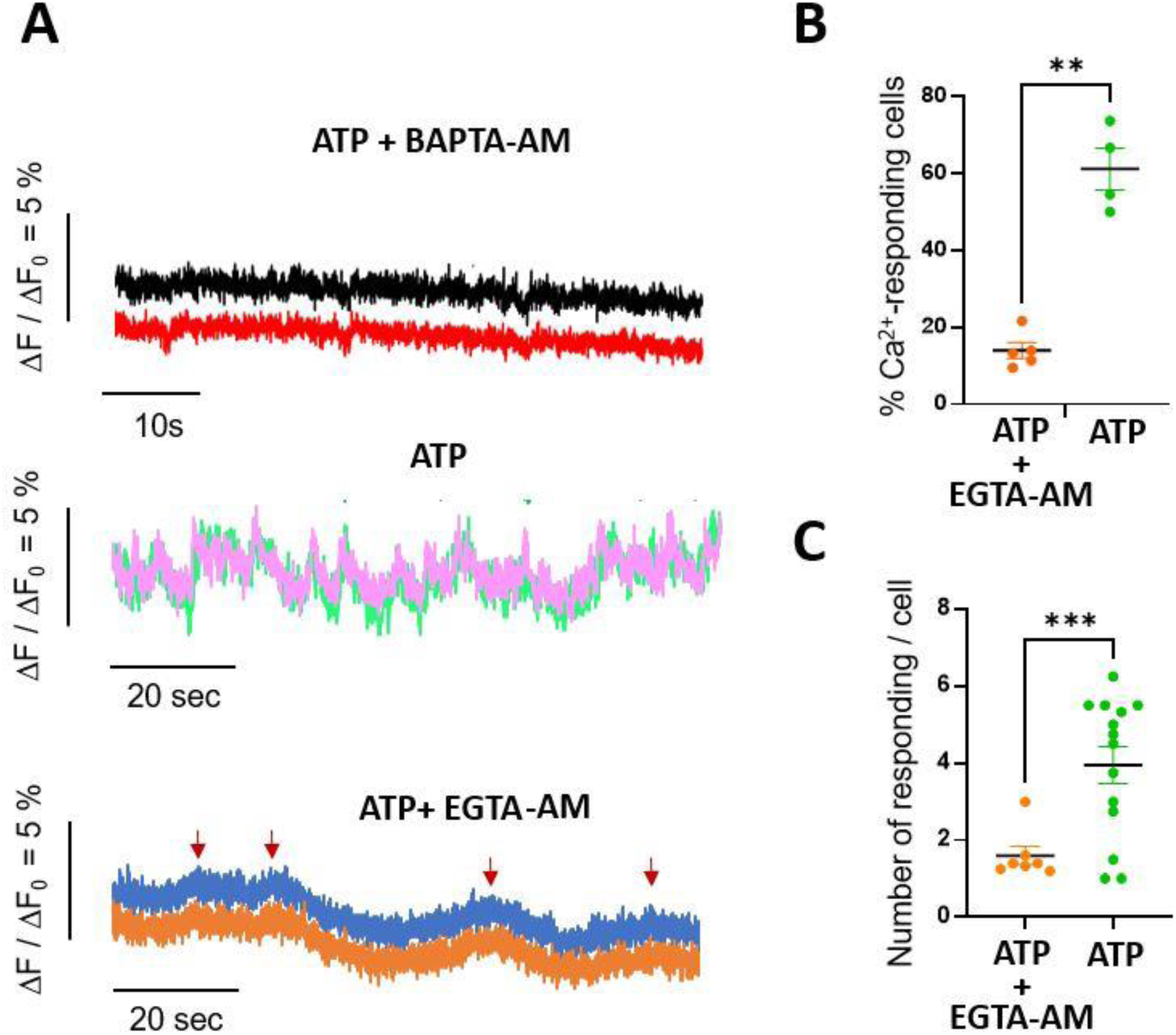
Intracellular Ca^2+^ chelation inhibits CCRICs. HeLa cells were loaded with the fluorescent indicator Cal-520 in the presence or absence of 20 μM EGTA-AM or BAPTA-AM. Samples were stimulated with 150 nM ATP and subjected to high speed Ca^2+^ imaging at a frequency of one acquisition every 22 ms. **A**, Representative traces of Ca^2+^ variations in 2 subcellular regions of the same cell in samples treated in the presence or absence of the indicated inhibitor. No Ca^2+^ responses were observed for cells treated with BAPTA-AM (N = 3, > 65 cells). EGTA-AM treatment led to an inhibition of Ca^2+^ responses, associated with small variations in the Ca^2+^ baseline that were arbitrarily scored as flattened Ca^2+^ pseudo-responses (ATP+EGTA-AM, red arrows). **B**, Percent of cells exhibiting Ca^2+^ responses (N > 3, cells > 62). **C,** average number of responses per cell. (N > 3, n > 62). Mann-Whitney test. **: p < 0.01; ***: p < 0.001.

**Fig. S8.**
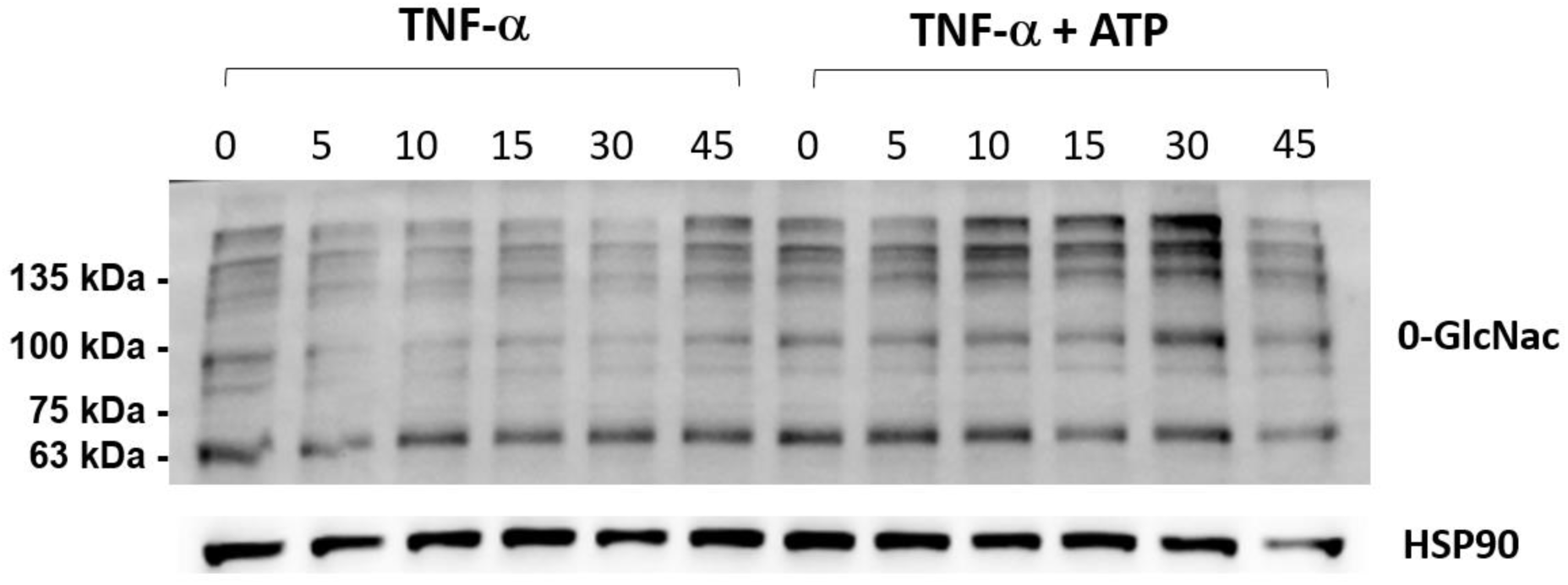
Effects of ATP on TNF-α-induced profiles of O-GlcNacylation in HeLa cells. HeLa cells were stimulated with 10 ng/ml TNF-α alone or co-stimulated with 10 ng/ml TNF-α and 150 nM ATP. Representative blots of cell lysates analyzed by Western blot at the time points indicated in minutes using the indicated antibodies.

